# Csb1 moonlighting gives rise to functional redundancy with Csb2 in processing the pre-CRISPR transcript in type I-G CRISPR-Cas system

**DOI:** 10.1101/2022.08.19.504415

**Authors:** Sunanda Chhetry, B Anand

## Abstract

Archaea and bacteria use CRISPR-based adaptive immunity to limit the genome invasion by phages. Among the type-I CRISPR variant, the newly discovered type I-G exhibits unusual variation in the composition and architecture of Cas proteins. In order to understand how these structural differences, contribute to functional adaptations, we probed how the maturation of CRISPR RNA differs with respect to other well studied type I CRISPR variants. Type I-G consists of three Cas proteins, viz, Csb1, Csb2 and Csb3 that are predicted to form the ribonucleoprotein surveillance effector complex. We show that Csb2 from *Bifidobacterium animalis* is a metal independent endonuclease that cleaves site-specifically within the 5’ region of the CRISPR repeat RNA. The catalytic activity resides within the C-terminal region that is homologous to Cas6. Interestingly, Csb2 processes the pre-CRISPR transcript both as a stand-alone enzyme and as a subunit of the Cascade/I-G complex that comprises of Csb1, Csb2 and Csb3 in association with crRNA. Surprisingly, we discovered that Csb1-which is homologous to Cas7 that is catalytically inert in other type I systems-also shows metal independent RNase activity that is functionally analogous to Csb2 in processing the pre-CRISPR RNA. The presence of dual nucleases in the Cascade/I-G complex enhances the efficiency of CRISPR-based immunity. We suggest that the Csb1 moonlighting engenders functional redundancy between Csb1 and Csb2 that in turn could compensate for the intrinsic instability of Csb2 and accelerate the maturation of crRNA.

## Introduction

Archaea and bacteria employ the CRISPR-Cas system as an adaptive immune mechanism to defend themselves from foreign mobile genetic elements (MGE) (1–3). CRISPR consists of an array of short conserved DNA-repeat sequences (~30-40 bp) that are interspaced by stretches of variable sequences called spacers. This array of repeats and spacers typically lie in close proximity with associated genes that encode for Cas proteins. The spacer sequences are derived from the invading MGE and integrated into the CRISPR array (4–7). The defense mechanism of the CRISPR-Cas system comprises of three operational stages namely (i) adaptation, (ii) maturation, and (iii) interference. CRISPR adaptation involves the recognition of foreign MGE followed by its subsequent polarized integration into the leader proximal region of the CRISPR array. CRISPR maturation follows next, where the CRISPR array is transcribed and processed to yield mature CRISPR RNA (crRNA). On this crRNA, either a multiple (Class 1) or a single Cas protein(s) (Class 2) co-assemble(s) to form a ribonucleoprotein (RNP) effector complex (8, 9). In the final step, the RNP complex guided by the crRNA recognizes the target and facilitates its cleavage, thus protecting the host. Based on the repertoire of *cas* genes and sequence similarity between the Cas proteins and locus architecture, the CRISPR-Cas system is divided into two major classes namely, Class1 and Class2, which are further divided into six types (type I-VI) and several subtypes. Type I, III, and IV have multisubunit effector complexes, whereas, type II, V, and VI possess single subunit effector complexes (8, 9).

The processing of the repeat-spacer transcript into mature crRNA is one of the important steps in the CRISPR defense. In the Class 2, maturation of crRNA is achieved by RNase III in association with Cas9 (type II), Cas12 (type V), and Cas13 (type VI), respectively (9–12). Whereas in the Class 1, Cas6 (most variants in type I, type III-C, D, E, and F and type IV) and Cas5 (type I-C) process the pre-CRISPR transcript (13–20). Cas6 belongs to the RAMP superfamily and it is a metal independent endoribonuclease. It consists of an RNA recognition motif (RRM) domain and a Glycine-rich loop that helps in binding to the premature crRNA (13, 16, 17, 21). The RNase follows an acid-base hydrolysis mechanism that produces a 2’-3’ cyclic intermediate, finally yielding products with 5’-OH and 3’-P ends (22–24). Strikingly in type I-C system, where Cas6 is absent, Cas5 assumes its role (14, 15, 18). So, in type I system, Cas6 and exceptionally Cas5 play an important role in crRNA maturation.

Type I-G is a newly identified atypical CRISPR-Cas system that shows tremendous variations in domain rearrangements (8, 9). One amongst these is the fusion of *cas5* and *cas6* to form *csb2*. The rest of the effector module includes Csb1 (Cas7 homolog) and Csb3 (Cas8 homolog). Though both Cas5 and Cas6 are separately shown to be essential for crRNA maturation in other type I CRISPR variants, how this fusion influences the processing of pre-CRISPR transcript, assembly and structural architecture of the RNP effector complex, and CRISPR based nucleic acid targeting in type I-G is not clear. In this study, we investigated the mechanism of crRNA maturation in type I-G system by probing the functional mechanism of Cas proteins from *Bifidobacterium animalis subsp. lactis*. We report that Csb2 is an RNA specific endonuclease that cleaves the pre-crRNA at a specific position from the 5’ end. Unlike in other type I systems, Csb1/I-G – the Cas7 homolog – also shows RNase activity that is comparable to Csb2, thus exhibiting functional redundancy in processing the pre-crRNA.

## Results

### Csb2 is an RNase that site-specifically cleaves the CRISPR repeat RNA

In *B. animalis*, we observed that the effector module in type I-G system consists of three Cas proteins, viz, Csb1, Csb2 and Csb3 (Figure 1A). We performed BLAST analysis on these proteins for the presence of conserved domains. We found that Csb1 is related to Cas7, Csb2 shows similarity to both Cas5 and Cas6 domains and Csb3 is related to Cas8. Typically, in type I systems, Cas5 or Cas6 is known to facilitate the maturation of CRISPR RNA. Therefore, we decided to biochemically investigate the role of Csb2 in CRISPR RNA maturation. We expressed *csb2* from *B. animalis* in *E*. *coli* BL21(DE3) and purified the protein to homogeneity (Figure S1A). It is worth mentioning that Csb2 is unstable and gets degraded within two weeks even at −80 °C. Therefore, all the assays were performed either immediately or within a couple of days from the purification (vide methods). To test its nuclease activity, we designed a 5’ 6-FAM labelled 36 nucleotide (nt) CRISPR repeat RNA from *B. animalis*, hereafter referred to as RNA_WT_ (Figure 1B). When Csb2 was incubated with RNA_WT_, we noted a prominent small RNA fragment, suggesting that Csb2 is an RNA specific endonuclease (Figure 1C, S1B). However, even after prolonged incubation, no further processing was observed, suggesting that Csb2 cleaves at a specific region within the repeat RNA (Figure S1B). In order to identify the cleavage site within RNA_WT_, we performed alkaline hydrolysis and RNase T1 based structural mapping. This revealed that Csb2 cleaves the repeat RNA at 5 nt from the 5’ end, that is between C_5_ and A_6_ (Figure 1D). Both Cas5 and Cas6 cleave the repeat RNA generating fragments that comprise of 5’ hydroxyl (OH) and 2’,3’-cyclic phosphate RNA ends (13, 16, 22, 23). To understand the nature of the RNA ends generated by Csb2, we treated the cleavage product with T4 Polynucleotide Kinase (PNK) in an acidic condition such that it favors removal of phosphates. On analyzing the reactions on a denaturing PAGE, it was found that the T4 PNK treated fragments migrated slower than that of the untreated ones indicating the conversion of the phosphate group to a less negatively charged hydroxyl group (Figure 1E). To further confirm this, we attempted to ligate the 3’ end of the dephosphorylated 5’ cleavage fragment to the phosphorylated 5’ end of the 3’ cleavage fragment using T4 RNA ligase. However, we could not succeed (lane 8 in Figure 1E) as the size of the generated RNA fragment (5 nt) is too small for T4 RNA ligase to act upon (25). Thus, based on the difference in migration, it appears that RNA cleavage by Csb2 yields two types of fragments: (i) one which has a 5’ phosphate and 2’,3’-cyclic phosphate (5’ fragment) and (ii) the other where both 5’ and 3’ ends have hydroxyl-group (3’ fragment). Further we wanted to test whether Csb2 possesses any DNase activity. Towards this, we incubated Csb2 with circular and linear DNA substrates with or without metal. (Figure S1C - F). While we observed nickase activity on circular DNA (dsDNA) at high concentration, there was no appreciable nuclease activity on linear dsDNA. This suggests that Csb2 is an RNA specific endonuclease that cleaves the CRISPR repeat RNA at a specific site.

**Figure 1.**
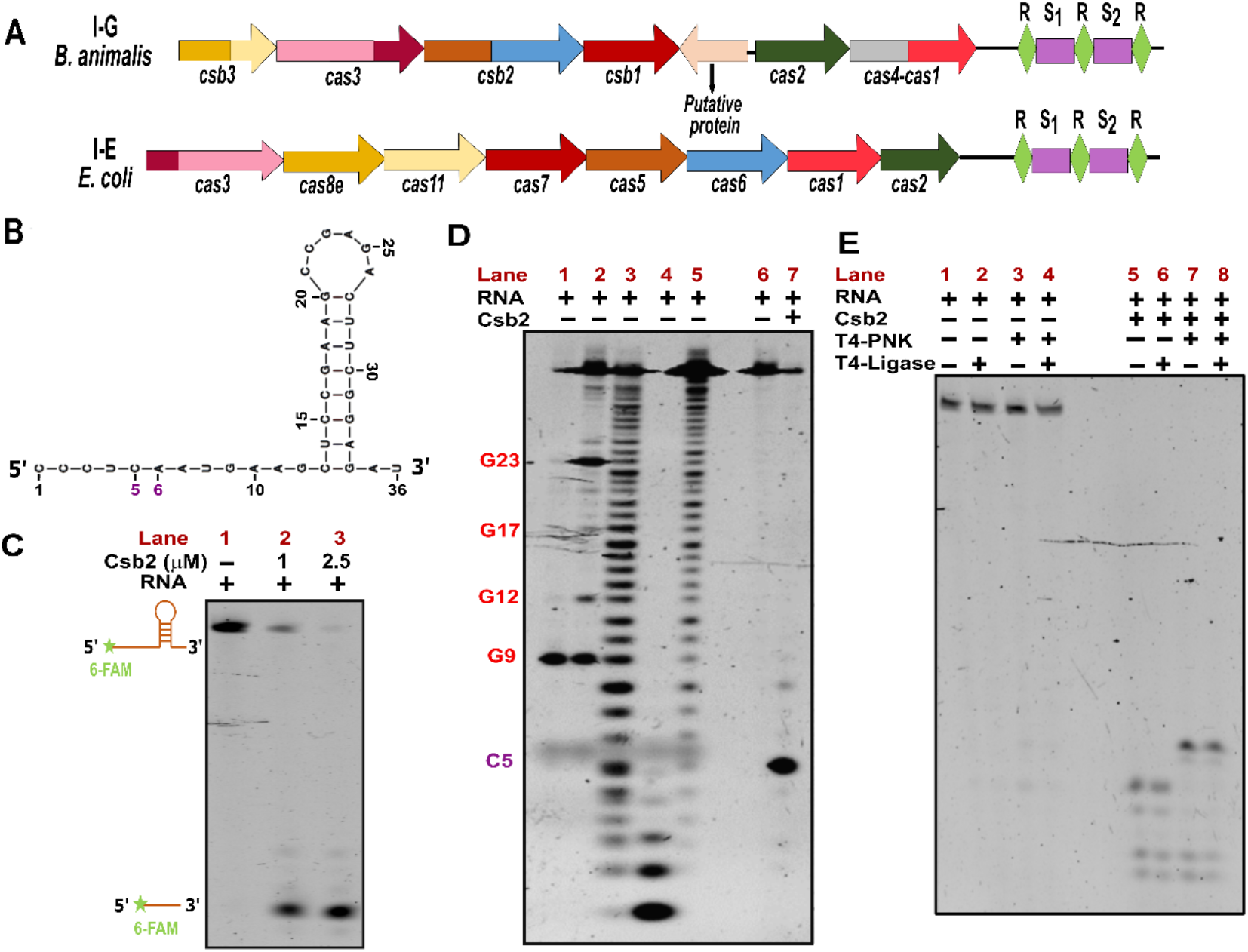
Csb2 is an RNase. **(A)** A schematic representation of the CRISPR-Cas locus of the well characterized type I-E from *E. coli* and type I-G from *B. animalis*. Genes are drawn to represent the locus architecture of the respective subtype. Repeats (R) are represented as green diamonds, whereas, the spacers (S) are shown as rectangles in magenta. **(B)** Predicted secondary structure representation of the *B. animalis* CRISPR repeat RNA is shown. The secondary structure prediction was made using the MFOLD Web Server (26). The 5^th^ and the 6^th^ residues from the 5’ end are indicated. **(C)** A 20% denaturing PAGE for assessing the RNase activity of Csb2 (1 μM and 2.5 μM) on 5’ 6-FAM labelled repeat RNA substrate is shown. The schema on the left shows both the substrate and the cleaved fragment. **(D)** A 20% denaturing PAGE is shown for mapping the cleavage site using an RNase T1 digestion ladder (lane 1 and 2) and an alkaline hydrolysis ladder (lane 3, 4 and 5). The RNase T1 digestion ladder in lane 1 and 2 was generated by treating the 5’ 6-FAM labelled repeat RNA with 1 and 0.1 units of RNase T1 enzyme, respectively. The alkaline hydrolysis ladder in lane 3, 4 and 5 was generated by varying the hydrolysis incubation times (5 and 10 min at 95 °C and 20 min at 60 °C), respectively. Positions that are specific to cleavage by RNase T1 are indicated as G9 through G23 in red. The cleavage site was mapped between C5 and A6, from the 5’ end. **(E)** A 20% denaturing PAGE is displayed for deciphering the nature of cleaved RNA fragment by Csb2. The cleaved RNA product was treated with T4 PNK at pH 5.2 in the absence of ATP to replace terminal phosphates with hydroxyl groups. The difference in migration between T4 PNK treated and untreated samples is attributed to the charge difference between these groups.

### The upstream sequence of the RNA stem is crucial for the nuclease activity

MFOLD prediction of repeat RNA suggests that it adopts a structured RNA, which is typical for several type I systems (26). To further investigate the sequence and structural preference of Csb2 for the RNA substrate, we designed several RNA constructs to generate structural and sequence variations in the substrate RNA (Supplementary table S2, Figure S2). Upon incubating Csb2 with each variant of RNA_WT_ that has altered stem-loop structure, we observed cleavage, suggesting that the stem-loop is not essential for the nuclease activity (Figure 2A-B). To assess the sequence conservation, we made sequence alterations upstream and downstream of the cleavage site. Most of the mutations do not alter the catalytic activity. However, we observed that deleting the 5’ region upstream of the stem compromises the RNA processing (Figure 2A-B), thus confirming the importance of this 5’ region for the nuclease activity.

**Figure 2.**
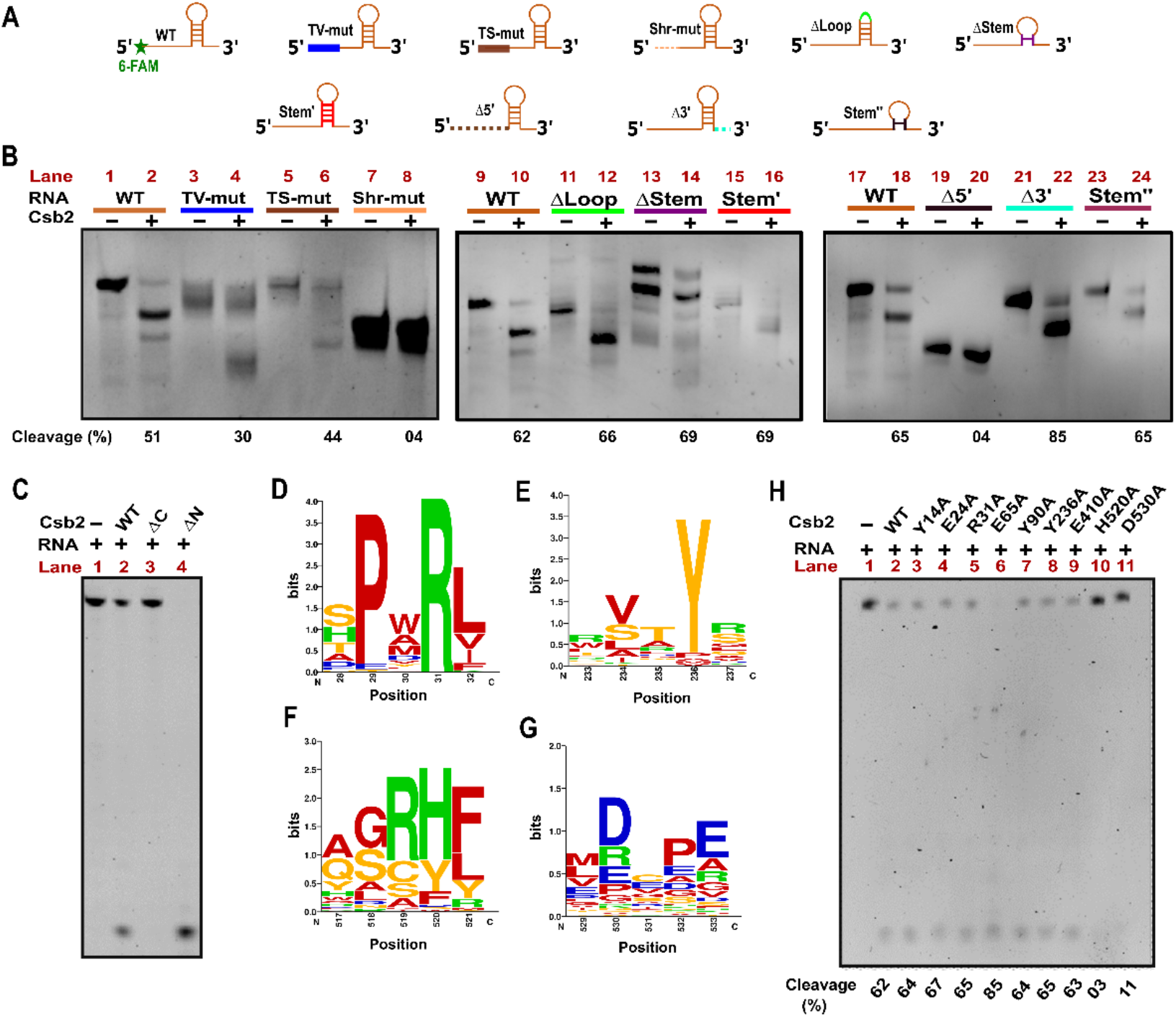
RNase activity of Csb2. (**A**) Schematic representation of the various RNA substrates used for characterizing substrate specificity of Csb2 is presented. The predicted secondary structure and the nomenclature of the respective RNA mutants are described in Figure S2 and Table S2. The variants of the wildtype (WT) substrate are indicated for the respective mutations as transversion (blue), transition (brown), deletion (dashed lines in green, purple, and black), and inversion (red). (**B**) An EtBr stained 20% denaturing PAGE is shown for assessing RNA substrate specificity of Csb2 (1 μM) on structural and sequence variants of RNA_WT_ (0.5 μM) as described in (A). The percentage of cleaved RNA fragment is indicated below the gel. (**C**) A 20% denaturing PAGE shows the RNase activity of 1 μM of truncated version of Csb2 such as the ΔC (lane 3) and ΔN (lane 4) on 0.5 μM 5’ 6-FAM labelled repeat RNA substrate, RNA_WT_, in comparison to the WT Csb2 (lane 2). (**D-G**) The sequence logo depicts the conservation profile of Csb2 orthologs (from 57 organisms). The X-axis indicates the amino acid residue number, whereas, the Y-axis indicates the extent of its conservation (in bits). (**H**) A 20% denaturing PAGE shows the RNase activity of 1 μM of the Csb2 point mutants Y14A, E24A, R31A, E65A, Y90A, Y236A, E410A, H520A, and D530A (lanes 3-11, respectively) on 0.5 μM of 5’ 6-FAM labelled repeat RNA substrate, in comparison to WT (lane 2). The percentage of cleaved RNA fragment is indicated below the gel.

### The C-terminal Cas6-like domain harbours the catalytic activity

Having confirmed that Csb2 is an RNase, we intended to probe which region of the Csb2 harbours the nuclease activity. To resolve this, we made domain assignments using the FFAS server (27) that suggested that the N-terminal region (1-250 aa) is homologous to Cas5 and the C-terminal region (251-545 aa) is homologous to Cas6 (Figure S3A). In order to assess whether Cas5-like or Cas6-like (or both) domain harbours the catalytic activity, we made N (Δ*N*) and C (Δ*C*) terminal deletions. This showed that Δ*N* retains the nuclease activity whereas the Δ*C* is inert (Figure 2C). In line with WT, Δ*N* and Δ*C* were inept in processing the Shr-mut RNA (Figure S3B). This suggests that the C-terminal Cas6-like domain is catalytic whereas the N-terminal Cas5-like domain is inert in type I-G system.

After identifying the Catalytic domain in Csb2, we proceeded to identify the active site residues. We inspected the orthologs of Csb2 across 57 organisms by performing a multiple sequence alignment using Clustal Omega (28). Among the conserved residues, we selected the charged residues (Figure 2D-G and S6) for site directed mutagenesis (SDM). The selected residues were substituted with alanine (Y14A, E24A, R31A, E65A, Y90A, Y236A, E410A, H520A, and D530A). All the mutants were cloned, confirmed by sequencing and the respective proteins were purified for assessing the nuclease activity. This showed that except H520A and D530A, the rest of the mutants retained the RNase activity similar to WT (Figure 2H). This suggests that H520 and D530 are the catalytic residues in Csb2. Notably, both H520 and D530 are situated in the C-terminal region of Csb2, thus lending further support to the catalytic role of C-terminal Cas6-like domain.

### Dual nucleases in the Cascade effector complex

Having characterized the nuclease activity of Csb2, we proceeded to address whether Csb2 processes the pre-CRISPR transcript either alone or as part of the effector complex. In other type I systems, Cas proteins such as Cas5, Cas6, Cas7 and Cas8 typically form the core of the Cascade complex in association with crRNA. We predicted that Csb1 is homologous to Cas7, Csb2 harbours N-terminal Cas5-like and C-terminal Cas6-like domains and Csb3 is homologous to Cas8. Therefore, these proteins were identified as potential constituents of the Cascade complex in type I-G system. To test whether these proteins assemble to form the Cascade complex, we cloned *csb1*, *csb2*, and *csb3* from *B. animalis* as a polycistronic construct (pCascade/I-G) and co-expressed with the CRISPR repeat-spacer array (pCR_Array/I-G) (Figure 3A, vide methods). Indeed, we could identify these proteins while analyzing the purified complex on SDS PAGE, suggesting that Csb1, Csb2, and Csb3 form a complex with crRNA (Figure 3B). The nucleic acid isolated from the purified complex was treated with DNase. After the DNase treatment, a prominent band was noted whose size corresponded to the predicted length (approximately 60-70 nt) of crRNA when analysed on a denaturing PAGE (Figure S4A). This lends further support that Csb1, Csb2, and Csb3 indeed co-assemble to form a complex with crRNA. Motivated by this, we asked whether the effector complex is capable of processing the CRISPR repeat RNA. Before performing the nuclease assay, we passed the RNP effector complex through a 100 kDa membrane filter to ensure that the effector complex is intact (vide methods). We noted that the effector complex showed endonuclease activity when incubated with CRISPR repeat RNA and that was more efficient than the Csb2 nuclease activity (Figure 3C). This suggests that the pre-CRISPR transcript is processed by Csb2 as a stand-alone protein as well as a subunit of the RNP effector complex. To validate that the nuclease activity of the effector complex is due to Csb2, we reconstituted the complex *in vivo* (vide methods) with Csb2 nuclease-dead mutant (H520A). Intriguingly, this mutant Cascade complex showed nuclease activity similar to WT Cascade complex, suggesting that there exists an additional nuclease within the complex (Figure 3C).

**Figure 3.**
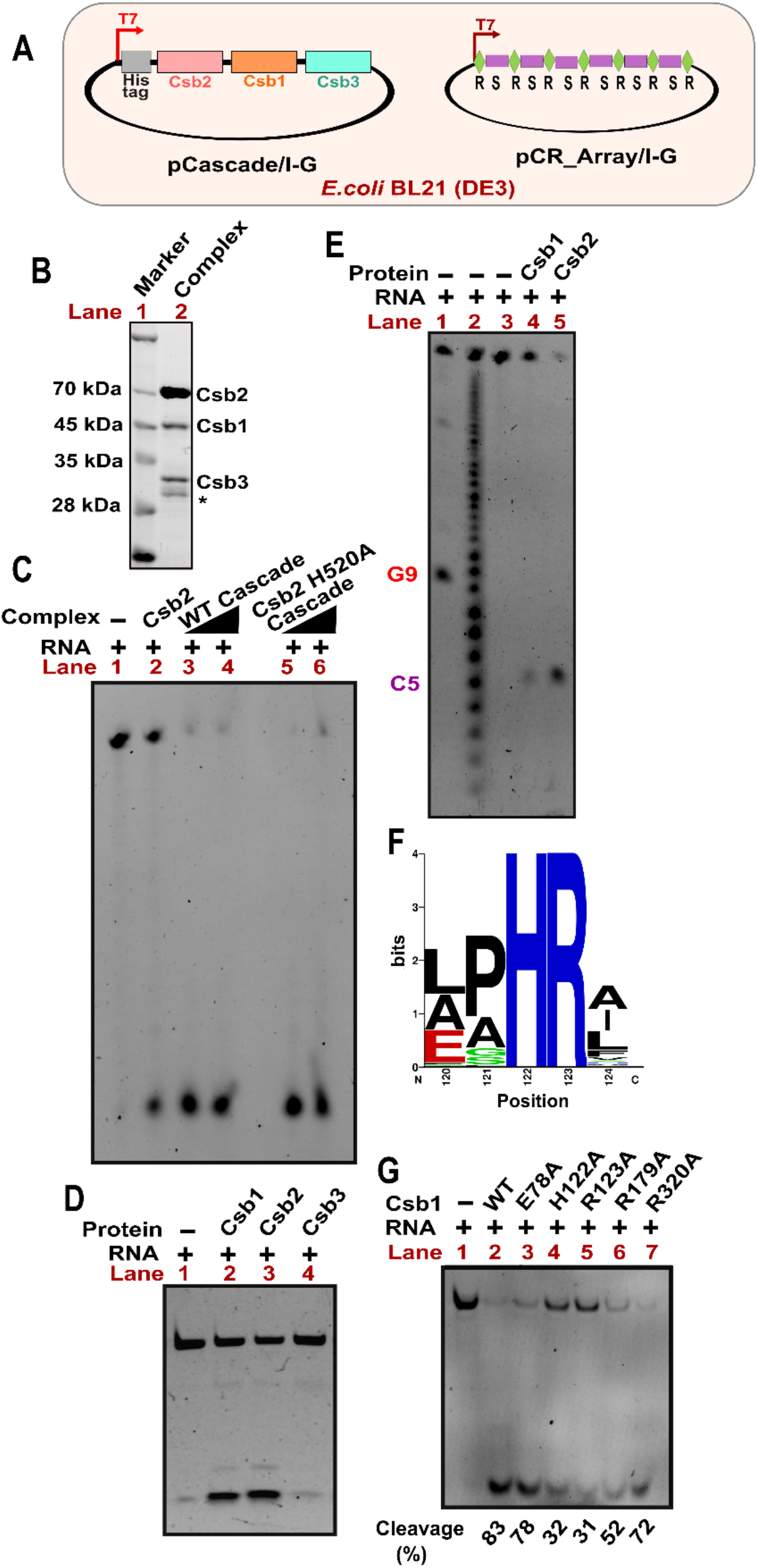
Csb1 is an RNase that is functionally redundant to Csb2. (**A**) A schematic representation of the constructs used for overexpression and *in vivo* reconstitution of the cascade complex is presented. Cascade/I-G (pCascade/I-G) and crRNA (pCR_Array/I-G) were co-expressed upon IPTG induction in *E. coli* BL21 (DE3) (**B**) A 10% SDS PAGE shows the individual subunits of the purified Cascade complex (lane 2). A protein marker is loaded on lane 2 to validate the molecular sizes. The respective subunits, namely, Csb1, Csb2 and Csb3 are also indicated on the right. The band indicated as * depicts progressive degradation of Csb2. (**C**) A 20% denaturing PAGE shows the RNase activity of 1 μM of Cascade complex (lane 3, 4) as well as Cascade (Csb2 H520A) complex (lane 5, 6) on 0.5 μM of 5’ 6-FAM labelled repeat RNA substrate. The Csb2 that was incubated with RNA substrate is shown in lane 2 as control. (**D**) A 20% denaturing PAGE shows the RNase activity of Csb1 (lane 2), Csb2 (lane 3), and Csb3 (lane 4) on 5’ 6-FAM labelled repeat RNA substrate. (**E**) A 20% denaturing PAGE is shown for mapping the cleaved RNA fragment by Csb1(lane 4) using an RNase T1 digestion ladder (lane 1) and an alkaline hydrolysis ladder (lane 2). The RNA substrate that was incubated with Csb2 separately in lane 5 is also indicated. Positions specific to cleavage by RNase T1 are indicated as G9 in red. The cleaved RNA fragment indicated as C5 (in purple) maps to the 5^th^ nucleotide from the 5’ end of the repeat RNA between C_5_ and A_6_. (**F**) The sequence logo depicts the conservation profile of amino acids in Csb1 across the type I-G system (57 organisms). The X and Y-axis indicate the residue number and the extent of its conservation (in bits), respectively. (**G**) A 20% denaturing PAGE shows the RNase activity of the Csb1 point mutants E78A, H122A, R123A, R179A, and R320A (1 μM each) (lanes 3-7), respectively, on 0.5 μM of 5’ 6-FAM labelled repeat RNA substrate, in comparison to WT (lane 2). The percentage of cleaved RNA fragment is indicated below the gel.

### Csb1 is an RNase that is functionally redundant to Csb2

The mutant Cascade complex (Csb2 H520A) retaining the RNase activity (Figure 3C) prompted us to hypothesize the presence of another RNase within the Cascade complex. To test this possibility, we proceeded to investigate the nuclease activity of Csb1 and Csb3. We individually cloned, expressed and purified these proteins and tested for the nuclease activity (Figure S4B-C). This showed that Csb3 is inert whereas Csb1 is an RNase that cleaves RNA_WT_ between C_5_ and A_6_ even in the absence of divalent metal ion (Figure 3D & E). Analysis of the cleaved products suggested that the cleavage has generated one fragment with 5’ phosphate and 2’,3’-cyclic phosphate (5’ fragment) and the other with a 5’ and 3’ hydroxyl-group (3’ fragment) (Figure S4D). These suggest that the preference for cleavage site and the nature of the cleaved fragments are very similar to the nuclease activity of Csb2. To confirm that the nuclease activity emanates from Csb1, we made alanine scanning mutagenesis (E78A, H122A, R123A, R179A and R320A) of conserved amino acids (Figure 3F, S4E-G & S5) and tested for the RNase activity. This showed that H122A and R123A exhibited reduced RNase activity compared to other mutants, suggesting that H122 and R123 are the catalytic residues of Csb1 (Figure 3G). To assess the sequence specificity against CRISPR repeat RNA, we treated Csb1 with different RNA substrates that show variation in structure and/or sequence. While Csb1 showed no structural or sequence specificity for its substrate in the stem or the loop region of the RNA, deleting the 5’ region upstream of the stem (Shr-mut and Δ5’) significantly compromised the RNA substrate processing, whereas a minor reduction was observed when the 5’ region was transversionally mutated or the stem regions were interchanged (Figure S4H). This suggests that the 5’ region of the CRISPR repeat RNA is important for the RNase activity of Csb1. The close functional overlap between Csb1 and Csb2 towards the CRISPR repeat RNA processing suggests that the type I-G system exhibits functional redundancy in processing the pre-CRISPR transcript.

### Functional redundancy accelerates crRNA maturation

After identifying Csb1 as an RNase, we proceeded to test whether the nuclease activity observed in Cascade (Csb2 H520A) complex is due to Csb1. To address this, we produced single (Csb1 H122A and R123A) and double (Csb1 H122A_Csb2 H520A and Csb1 R123A_Csb2 H520A) mutants of Csb1 and Csb2 and reconstituted the mutant Cascade complex *in vivo*. After testing the nuclease activity, we found that only the double mutants, viz, Csb1 H122A_Csb2 H520A and Csb1 R123A_Csb2 H520A were inert whereas the single mutants (Csb1 H122A and R123A) retained the nuclease activity albeit at a low level (Figure 4A). This suggests that the loss of Csb1 nuclease activity is compensated by Csb2 within the Cascade complex. This further lends support that Csb1 and Csb2 act as dual nucleases within the Cascade complex. To understand the consequences of this functional redundancy on CRISPR interference, we performed the CRISPR interference assay *in vivo* using *E. coli* as a surrogate host. To functionally reconstitute the type I-G CRISPR machinery in *E. coli*, we introduced plasmid-borne copies of the Cascade (pCascade), Cas3 (pCas3), and target (pT) or, non-target (pNT) into *E. coli* IG-CR strain (Figure 4B, vide methods). This strain expresses the pre-CRISPR transcript from *B. animalis* from a chromosomally integrated cassette (vide methods). A fully functional CRISPR machinery comprising of Cascade complex and Cas3 reduced the transformation efficiency of pT (target) as opposed to the pNT (non-target), suggesting that the CRISPR-Cas system of type I-G is functional towards the target substrate (Figure 4C & S4I). Encouraged by this, we tested the effect of single and double mutants of Csb1 and Csb2 on the CRISPR interference (Figure 4D). We observed that the single mutants, viz, Csb1 H122A, Csb1 R123A, and Csb2 H520A showed enhanced transformation efficiency compared to WT, whereas the double mutants (Csb1 H122A_Csb2 H520A and Csb1 R123A_Csb2 H520A) showed better transformation efficiency than the single mutants (except Csb1 H122A), suggesting that the mutations in both Csb1 and Csb2 abrogates the CRISPR interference. Taken together, this further suggests that the presence of dual nucleases (Csb1 and Csb2) enhances the efficiency of CRISPR-based target elimination, possibly, by accelerating the processing of pre-CRISPR transcript.

**Figure 4.**
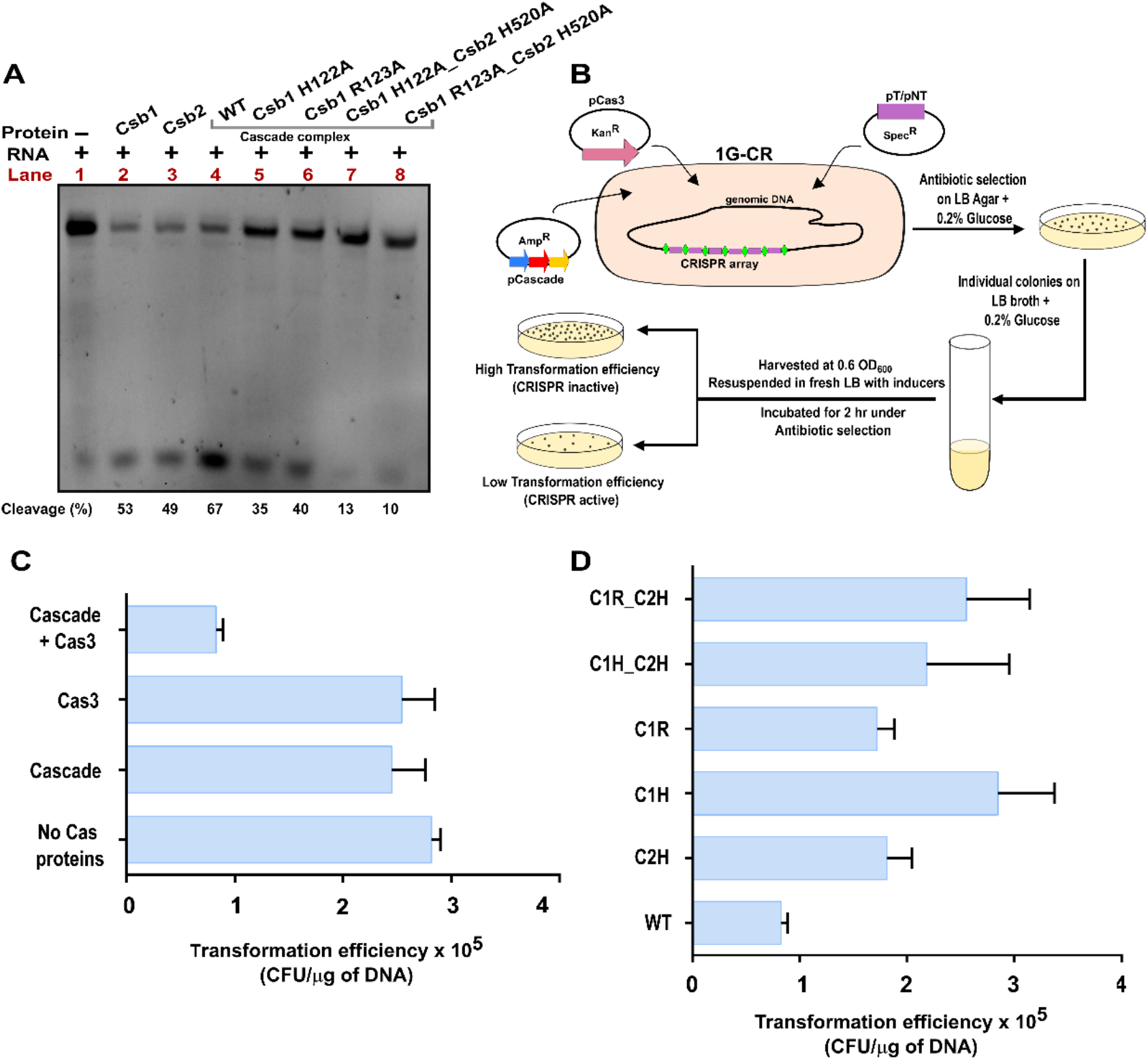
Dual nucleases participate in the maturation of crRNA in type I-G. (**A**) A 20% denaturing PAGE depicts the RNase activity of WT (lane 4) and mutant Cascade complexes (1 μM each) on 5’ 6-FAM labelled repeat RNA substrate, RNA_WT_ (0.5 μM). Mutants include Csb1 H122A (lane 5), Csb1 R123A (lane 6), Csb1 H122A_Csb2 H520A (lane 7), and Csb1 R123A_Csb2 H520A (lane 8). The RNA substrate was also incubated with Csb1 (lane 2) and Csb2 (lane 3) for a comparative analysis. The percentage of cleaved RNA fragment is indicated below the gel. (**B**) A schematic representation of the *in vivo* interference assay is shown. An *E. coli* IG-CR strain harbouring the CRISPR array from *B. animalis* was used as a surrogate host. Cascade/I-G complex (including its varied mutants) and Cas3 were co-expressed through IPTG inducible vectors harbouring ampicillin and kanamycin antibiotic resistance markers, respectively. The target sequence was inserted in 13S-R vector (T), whereas an empty 13S-R vector was used for the non-target plasmid (NT). Low transformation efficiency indicates a functional CRISPR interference that may be attributed to the successful maturation of crRNA. (**C**) Transformation efficiency is shown for *E. coli* IG-CR in response to the presence of the target plasmid and other components of the CRISPR interference machinery such as Cascade complex and Cas3. The error bars represent standard deviation from three independent trials. (**D**) Transformation efficiency of *E. coli* IG-CR was tested as in (C) in the presence of varied mutant forms of Cascade complex such as C2H (Csb2 H520A Cascade), C1H (Csb1 H122A Cascade), C1R (Csb1 R123A Cascade), C1H_C2H (Csb1 H122A, Csb2 H520A Cascade), C1R_C2H (Csb1 R123A, Csb2 H520A Cascade). The error bars represent the standard deviation observed across three independent trials

## Discussion

pre-CRISPR transcript processing is a cardinal step that ensures the maturation of crRNA that in turn guides the recognition of target DNA/RNA for successful CRISPR-Cas based genome defence. In most type I systems (except type I-C) studied so far, the mandate to produce the guide crRNA is fulfilled by Cas6, which processes the pre-CRISPR transcript by cleaving within the repeat region to liberate the individual guide crRNA (13, 16, 17, 21, 23). Cas6 is an endoribonuclease that typically consists of two ferredoxin-like (also referred as RNA recognition motif (RRM)) domains arranged in tandem and a glycine-rich region at the C-terminus (29). Exceptionally, in type I-C, the absence of Cas6 is compensated by a catalytically active Cas5, which is usually inert in other type I systems (14, 15, 18). Like Cas6, Cas5 also possesses a ferredoxin-like domain. Therefore, the fusion of Cas5-like (N-terminus) and Cas6-like (C-terminus) domains in Csb2 is intriguing as to which domain is catalytically active. We found that the nuclease activity resides at the C-terminal Cas6-like domain, whereas the N-terminal Cas5-like domain is catalytically inert (Figure 2C). Here, the nature of the Csb2-produced RNA fragments (the production of 5’-OH and 2’,3’ cyclic phosphate) are characteristic of both Cas5 and Cas6 nuclease activities (Figure 1E). However, unlike Cas6 (and Cas5) in other type I systems that show preference for the base of the stem-loop structure of the CRISPR repeats (13–16, 30–32), Csb2 shows no specificity towards the stem region, as altering the sequence, size and position of the stem do not affect the RNase activity (Figure 2B). However, it was observed that deleting the 5’ overhang of the repeat RNA abrogates the nuclease activity (Figure 2B), suggesting that the mode of specificity towards the repeat RNA is unconventional. Except in type I-E, where Cas6 processes the pre-CRISPR transcript as a subunit of the Cascade complex (24, 33)-Cas6 in other type I systems (A, B, C and F) and most type III systems (A-C) act as a stand-alone enzyme (15, 22, 23, 29, 30, 34–36). At variance with this, Csb2 in type I-G system exhibits the nuclease activity both as a stand-alone protein as well as a subunit of Cascade complex (Figure 3C). This suggests that the fusion of Cas5 and Cas6 in Csb2 has retained the characteristic nuclease activity of Cas6; however, it exhibits functional plasticity by showing similarity to both type I and III systems.

In type I-G, we discovered that the Cascade complex, in addition to Csb2, harbours another RNase –Csb1-that processes the pre-CRISPR transcript. Csb1 is homologous to Cas7; however, Cas7 in other type I systems (A-F) is generally inert and it forms the backbone of the Cascade complex (24, 33, 34, 37–48). In contrast, Cas7 in type III system cleaves the target RNA and this triggers the cyclic oligo adenylate (cOA) signalling leading to the indiscriminate cleavage of the host RNA (9, 49–53). Hitherto, no such precedence was noted in type I system. Therefore, Csb1 forms the founding member of catalytically active Cas7 in type I system. Intriguingly, though the domain architecture of Csb1 and Csb2 are not similar, they exhibit functional redundancy. In terms of the cleavage site on the CRISPR repeat, preference for 5’ overhang, and the nature of cleaved products, Csb1 is functionally very similar to Csb2 (Figure 3E and S4D, H). Yet, there are some subtle differences; Csb1 and Csb2 share ferredoxin domain and thus are structurally similar, yet, the catalytic residues in Csb1(H122 & R123) and Csb2 (H520 & D530) seem to be supplied from non-equivalent positions in the ferredoxin domain. This suggests that Csb1 has acquired the catalytic trait by convergent evolution that renders it functionally redundant with Csb2. What could be the evolutionary advantage of this functional redundancy? We noted that Csb2 is highly unstable and undergoes rapid degradation within few days even at low temperature. In order to elicit robust CRISPR-based defence, robust production of crRNA is crucial. This alludes to the possibility that Csb1 has evolved to be catalytic in type I-G to compensate for the instability of Csb2. Type III systems are reported to be the ancient CRISPR-Cas systems, from which the type I systems are proposed to have arisen (8). The fact that the Csb1 – a Cas7 homolog in type I – shows functional similarities to Cas7 in type III system suggests that type I-G could be the relic of the intermediate forms of the CRISPR-Cas system during the evolution of type I from type III system.

## MATERIALS AND METHODS

### Construction of bacterial strains and plasmids

The Genomic DNA of *Bifidobacterium animalis subsp lactis* (DSM 10140) was procured from DSMZ. The genes encoding Csb2/I-G and its point mutants such as Y14A, E24A, R31A, E65A, Y90A, Y236A, E410A, H520A and D530A as well as ΔN (1-250 aa) and ΔC (251-545 aa) truncation domains were PCR amplified and cloned into pET Strep II TEV LIC cloning vector (1R, Addgene #29664). The genes encoding Csb1/I-G, its point mutants, namely, E78A, H122A, R123A, R179A and R320A were PCR amplified and cloned into pET Strep II co-transformation cloning vector (13S-R, Addgene # 48328). Csb3/I-G was cloned into pQE2 (Qiagen) using Gibson assembly. A polycistronic construct consisting of genes encoding Csb2, Csb1 and Csb3 was designed and cloned into pQE2, and this will be hereafter referred as pCascade/I-G. Cascade complex with various point mutants like Csb2-H520A, Csb1-H122A, Csb1-R123A and a combination of these were similarly cloned into pQE2. A type I-G CRISPR array from *B. animalis* harboring six identical repeat-spacer units was placed under the control of a T7 promoter. This construct was commercially synthesized (GeneArt) and cloned into 13S-R (Addgene # 48328) to produce pCR_Array/I-G (Table S4). These constructs (pCR_Array/I-G and pCascade/I-G) were used to co-express and purify the *in vivo* assembled WT and mutant Cascade/I-G complexes. To construct the target plasmid (pT/I-G), target DNA sequence was cloned into 13S-R (Addgene # 48328). For testing the CRISPR interference *in vivo*, the WT CRISPR array (1.7 kb) from *B. animalis* including 175 nt upstream and 150 nt downstream, was integrated into the *attB* sites of the genome of *E. coli* IYK-12 using pOSIP-CT (Addgene #45981) to create the strain IG-CR via clonetegration (54). The gene encoding Cas3/I-G was cloned into pET Strep II TEV LIC cloning vector (1R, Addgene #29664). All the clones were confirmed by Sanger sequencing. The detailed information on all the constructs including primer sequences used in this study is mentioned in the Supplementary table S1, S3 and S4.

### Expression and purification of Cas proteins

In order to overexpress and purify Csb2 or its mutant variants, *E. coli* BL21 (DE3) cells were transformed with plasmids carrying the respective genes. These cells were grown in LB broth supplemented with 50 μg/ml kanamycin at 37 °C until OD_600_ reached 0.6 and then induced with 0.2 mM IPTG. The expression was continued at 25 °C for 12 hr and then the cells were harvested by centrifugation at 25 °C, 10,000 g for 5 min. The cell pellets were washed and resuspended in binding buffer A (20 mM Tris-Cl (pH 7.6), 500 mM NaCl and 6 mM β-mercaptoethanol) containing 1 mM phenylmethylsulfonyl fluoride (PMSF) and Protease inhibitor cocktail (Sigma-Aldrich) (20 mg/g of cell pellet). Cells were lysed by sonication (Pulse – 2 sec ON and 20 sec OFF) and clarified by centrifugation at 4°C and 20,000 g for 30 min. The supernatant was loaded onto a pre-equilibrated 5 ml StrepTrap HP (GE Healthcare) column. Upon binding, the column was washed with 10 CV of binding buffer A and then the protein was eluted in binding buffer A containing 2.5 mM d-Desthiobiotin (Sigma-Aldrich). The eluted fractions were further purified using a HiLoad 16/600 Superdex 75 prep grade column (GE Healthcare). The protein was concentrated using ultrafiltration through a 50 kDa cutoff filter (Sartorius), snap-frozen and stored at −80 °C until further use. We observed that Csb2 is highly unstable (instability index (II) (55) – 54.94) and undergoes spontaneous degradation within few days even at – 80 °C. Therefore, after the protein was purified, all assays with Csb2 were performed immediately or within a couple of days. Similarly, Csb1 and its mutant variants were purified and stored.

For purification of Csb3, *E. coli* BL21 (DE3) harboring pCsb3 was grown in LB broth supplemented with 100 μg/ml ampicillin at 37 °C until OD_600_ reached 0.6 and induced with 0.2 mM IPTG. The expression was continued at 25 °C for 12 hr and then the cells were harvested by centrifugation at 25 °C, 10,000 g for 5 min. The pellet was then resuspended in binding buffer B (20 mM Tris-Cl (pH 7.6), 300 mM NaCl, 30 mM imidazole and 6 mM β-mercaptoethanol) containing 1mM phenylmethylsulfonyl fluoride (PMSF) and Protease inhibitor cocktail (20 mg/g of cell pellet). After sonication (Pulse – 2 sec ON and 20 sec OFF), the lysate was centrifuged at 4 °C and 20,000 g for 30 min. The supernatant was loaded onto a pre-equilibrated 5ml HisTrap HP (GE Healthcare) column. After the sample binding, the column was washed with 10 CV of binding buffer B and eluted on a linear gradient of imidazole (30 mM to 1 M) in buffer B. The eluted fractions were loaded onto a pre-equilibrated HiLoad 16/600 Superdex 75 prep grade column (GE Healthcare) and purified further. The protein was concentrated using ultrafiltration through a 30 kDa cutoff filter (Sartorius), snap-frozen and stored at −80 °C until further used.

### *In vitro assay* for nuclease activity

To test the RNase activity of Csb1/I-G, Csb2/I-G and Csb3/I-G, a 36 nt RNA repeat sequence labelled with 5’ 6-FAM was chemically synthesized from IDT. 0.5 μM of this substrate was incubated with 1 μM of Csb1/I-G, Csb2/I-G and Csb3/I-G or their variants in a reaction buffer 1 containing 20 mM Tris-Cl (pH 7.6), 100 mM KCl and 6 mM β-Mercaptoethanol (β-ME) and incubated at 37 °C for 30 min. The samples were analyzed on a 20% (w/v) denaturing polyacrylamide gel and viewed in Bio-Rad gel documentation system.

Various mutants of the RNA substrate (Supplementary Table S1, S2 and Figure S2) were synthesized via *in vitro* transcription using T7 RNA polymerase, followed by DNase I treatment and phenol-chloroform extraction. They were further purified by gel extraction followed by ethanol precipitation. 0.5 μM of the gel-purified substrates were treated with 1 μM of Csb1/I-G and Csb2/I-G and the samples were analyzed on a 20% (w/v) denaturing polyacrylamide gel, stained with Ethidium Bromide (EtBr) and viewed in Bio-Rad gel documentation system. The percentage of RNA cleavage was quantified using densitometric analysis of gel images using Image lab package (Bio-Rad) and the values are indicated for the respective figures.

To assess the DNase activity of Csb2/I-G and Csb1/I-G, circular or linear pUC19 (dsDNA) was incubated with varied concentrations of Csb2/I-G in reaction buffer 1. The samples were visualized in 0.8% agarose gel using EtBr stain. The reaction buffer 1 were supplemented with Mg^2+^ ions, wherever necessary.

### Identification and characterization of cleavage site

An RNase T1 mediated digest was employed for analyzing the cleaved products (56). Briefly, for RNase T1 digestion reaction, CRISPR repeat RNA (5’ 6-FAM labelled) was treated with 0.1 units of RNase T1 in 10 mM Tris-Cl (pH 7), 100 mM KCl and 10 mM MgCl_2_. In order to size the generated fragments, an alkaline hydrolysis ladder was generated by heating the CRISPR repeat RNA (5’ 6-FAM labelled) with 1X alkaline hydrolysis buffer (50 mM Sodium carbonate (pH 9.2) and 1 mM EDTA) at 95 °C for 5 min and snap cooled in ice. These digests along with the test sample where the CRISPR repeat RNA (5’ 6-FAM labelled) was treated with Csb2/I-G and Csb1/I-G, respectively, were subsequently resolved on a 20% (w/v) denaturing polyacrylamide gel and visualized in a Bio-Rad gel documentation system.

To assess the nature of the fragment released by Csb2/I-G and Csb1/I-G activity on the repeat RNA, the cleaved fragment was isolated and treated with T4 polynucleotide kinase (T4-PNK) in 100 mM Tris-acetate (pH 6), 10 mM MgCl_2_ and 2 mM EDTA at 37°C for 30 min (57). The end repaired RNA was purified by ethanol precipitation and treated with 1 unit of T4 RNA ligase 1 in 50 mM Tris-HCl (pH 7.5), 10 mM MgCl_2_, 10 mM DTT, 1 mM ATP at 37 °C for 30 min, to ligate the 3’ end of the dephosphorylated 5’ cleavage fragment to the phosphorylated 5’ end of the 3’ cleavage fragment (25). It was subsequently resolved in 20% (w/v) denaturing polyacrylamide gel and visualized in a Bio-Rad gel documentation system.

### *In vivo* Cascade complex reconstitution

To reconstitute the Cascade complex *in vivo*, *E. coli* BL21 (DE3) was co-transformed with pCascade/I-G and pCR_Array/I-G. The cells were grown in LB broth in presence of 50 μg/ml kanamycin and 100 μg/ml spectinomycin until OD_600_ reached 0.6. Protein production was then induced by adding 0.2 mM IPTG and growth was continued at 16 °C for another 12 hours. The cells were harvested and resuspended in binding buffer C (20 mM Tris-Cl (pH 7.6), 30 mM NaCl, 10% glycerol and 6 mM β-mercaptoethanol) containing 1 mM phenylmethylsulfonyl fluoride (PMSF), Protease inhibitor cocktail (Sigma Aldrich) (20 mg/g of cell pellet) and RNase free – DNase I (NEB). Lysis was done in a cell disruptor (20 kpsi). The lysate was then clarified by centrifugation at 4 °C and 20,000 g for 30 min followed by an additional step of ultracentrifugation at 4 °C and 120,000 g for 2 h. The lysate was loaded onto a pre-equilibrated 5 ml HiTrap SP HP (GE Healthcare) column connected in series with 5 ml HiTrap Q HP (GE Healthcare) column. After binding, the column was washed individually with 10 CV of binding buffer C. The complex was eluted from the HiTrap Q HP column using a linear gradient of 30 mM to 1 M NaCl in buffer C. The eluted fractions were pooled and loaded onto a pre-equilibrated 5 ml HisTrap HP (GE Healthcare) column. After washing with 10 CV of buffer D (20 mM Tris-Cl (pH 7.6), 300 mM NaCl, 30 mM imidazole, 10% glycerol and 6 mM β-mercaptoethanol) the complex was eluted using a linear gradient of 30 mM to 1 M imidazole in buffer D. For the final step of purification, the fractions were loaded onto a pre-equilibrated HiLoad 16/600 Superdex 200 prep grade column (GE Healthcare) and eluted in the storage buffer (20 mM Tris-Cl (pH 7.6), 300 mM NaCl, 10% glycerol and 6 mM β-mercaptoethanol). The complex was concentrated using ultrafiltration through a 100 kDa cutoff filter (Merck) and analyzed on a 10% SDS PAGE for the presence of Cas subunits. Similar to Csb2, the Cascade complex was also observed to undergo spontaneous disintegration within few days even at −80 °C. Therefore, after the complex was purified, all assays were performed immediately or within a couple of days. To ascertain that no dissociated subunits were used while performing the experiments, each time the purified complex was re-passed through a 100 kDa cutoff filter and only the retentate was used for the *in vitro* experiments. The mutants of Cascade complex, namely pC1H_Cascade (Csb1 H122A), pC1R_Cascade (Csb1 R123A), pC2H_Cascade (Csb2 H520A), pC1H_C2H_Cascade (Csb1 H122A and Csb2 H520A) and pC1R_C2H_Cascade (Csb1 R123A and Csb2 H520A) were also reconstituted similarly (Supplementary Table S1 and S4). To isolate the crRNA from the intact Cascade complex, 500 μl of the concentrated sample was heated at 95 °C for 15 min and further subjected to phenol-chloroform extraction and ethanol precipitation. The sample thus obtained was treated with RNase-free DNase I (50 μg/ml) at 37 °C for 1 hr to remove any traces of DNA. The RNA that remained was visualized using a EtBr stained 20% denaturing polyacrylamide gel.

To assess the RNase activity of the Cascade complex, the 5’ 6-FAM labelled repeat RNA was incubated with WT Cascade complex or its variants in reaction buffer 1 containing 20 mM Tris-Cl (pH 7.6), 100 mM KCl and 6 mM β-Mercaptoethanol (β-ME) and incubated at 37 °C for 30 min. The samples were analyzed on a 20% (w/v) denaturing polyacrylamide gel and viewed in Bio-Rad gel documentation system.

### *In vivo* CRISPR interference assay

To establish a functional CRISPR interference assay system for type I-G *in vivo*, *E. coli* IG-CR cells harboring pCascade/I-G, pCas3/I-G, pT/I-G, and pNT/I-G in various combinations were used as surrogate host. As mentioned earlier, the type I-G CRISPR array was integrated into the chromosomal DNA of *E. coli* IYK12 to produce *E. coli* IG-CR using clonetegration. *E. coli* IG-CR was transformed with various combinations of constructs and plated onto an LB agar plate supplemented with the respective antibiotics (Supplementary Table S1, S3 and S4) and 0.2 % glucose to arrest any leaky expression. A single colony was picked to inoculate a fresh 5 ml LB broth containing the respective antibiotics (25 μg/ml chloramphenicol, 25 μg/ml kanamycin, 50 μg/ml ampicillin, 50 μg/ml spectinomycin) and 0.2% glucose and allowed to grow until OD_600_ reached 0.6 at 37 °C. 500 μl of the cells were centrifuged and the pellet was washed with fresh LB broth two times to remove any traces of glucose. The pellet was resuspended in 500 μl of fresh LB broth supplemented with the respective antibiotics as mentioned above and 0.02 mM IPTG and 0.2 % L-arabinose and allowed to grow at 180 rpm at 37 °C for about 2 hours. After this, the pellet was harvested and serially diluted upto 10^−5^ times and plated onto the LB agar plates supplemented with antibiotics and incubated at 37 °C for 10-12 hours. The number of colonies in each plate was counted, and the transformation efficiency was calculated using the following equation.

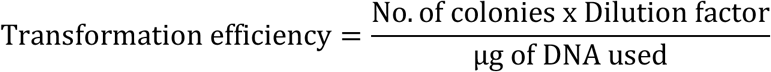

Transformation efficiency for various mutants of the Cascade complex (pC1H_Cascade, pC1R_Cascade, pC2H_Cascade, pC1H_C2H Cascade and pC1R_C2H Cascade) was calculated using a similar protocol.

## Acknowledgments

Vector pOSIP-CT (Addgene #45981) was a kind gift from Dr Drew Endy and Dr Keith Shearwin; p1R (Addgene #29664), and p13SR (Addgene #48328) were kindly provided by Dr Scott Gradia; E. coli IYB5101 was a kind gift from Dr Udi Qimron. We acknowledge the geniality of the aforementioned scientists for sharing their plasmids and bacterial strains. We acknowledge the funding from Science and Engineering Research Board [CRG/2019/003385] and Ignite Life Science Foundation [IGNITE/FG-OC/2021/005]. We thank Sudipta Mahapatra for her help with some experiments pertaining to Csb1, and all members of the MAB lab for their critical comments and suggestions.

## Competing Interest Statement

The authors declare no competing interests.

**Supplementary information** is available at BioRxiv online.

## Notes

### Competing Interest Statement

The authors have declared no competing interest.

