## Supplementary Information for "Csb1 moonlighting gives rise to functional redundancy with Csb2 in processing the pre-CRISPR transcript in type I-G CRISPR-Cas system"

Sunanda Chhetry and B. Anand*

Department of Biosciences and Bioengineering, Indian Institute of Technology Guwahati, Guwahati 781039, INDIA

*Author for Correspondence

### SUPPLEMENTARY INFORMATION

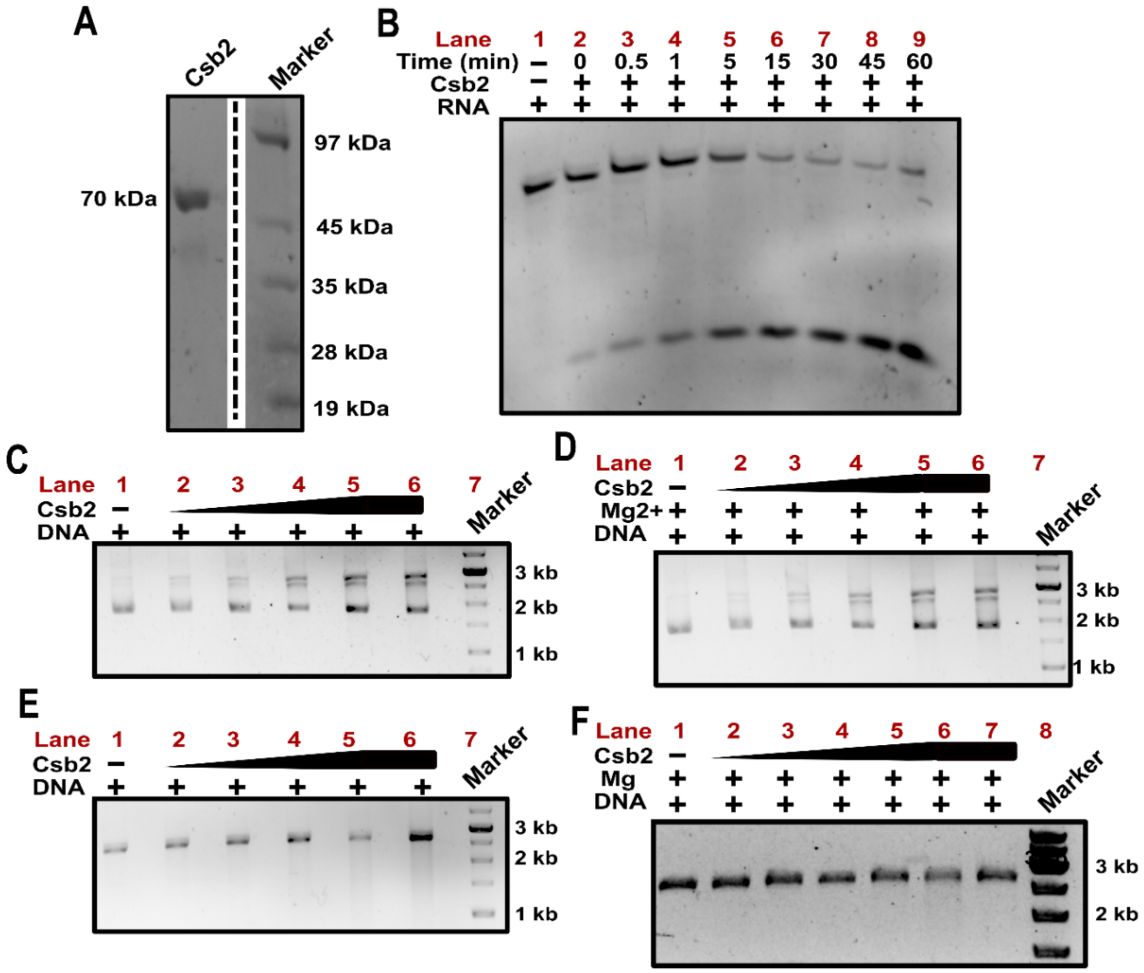


**Figure S1. (A)** A 15% SDS PAGE shows the purified 70 kDa Csb2 protein (marked as Csb2). A protein marker is loaded on right and the respective molecular weights are indicated. The vertical broken line indicates discontinuity between the lanes, which is introduced for the sake of clarity. (**B**) A 20% denaturing PAGE shows the time-dependent (0-60 min) RNase activity of Csb2 (1 μM) on 5' 6-FAM labelled WT repeat RNA substrate (0.5 μM). (**C, D**) A 1% agarose gel shows the DNase activity of increasing concentration of Csb2 (0.5, 1, 2.5, 5 μM) on circular dsDNA substrate in the absence (**C**) and presence (**D)** of divalent metal ion ($Mg^{2+}$), respectively (lanes 2-5). Lane 6 consists of DNA treated with 5 μM of Csb2, followed by Proteinase K treatment. A DNA marker is loaded in lane 7, and the respective bands are marked for reference. (**E**) A 1% agarose gel depicts the DNase activity of increasing concentration of Csb2 (0.5, 1, 2.5, 5 μM) on linear dsDNA substrate (lanes 2-5). Lane 6 consists of DNA treated with 5 μM of Csb2, followed by Proteinase K treatment. A DNA marker is loaded in lane 7, and the respective bands are marked for reference. **(F)** A 1% agarose gel shows the DNase activity of increasing concentration of Csb2 (0.5, 1, 2, 3, 5 μM) on linear dsDNA substrate (lanes 2-6) in the presence of divalent metal ion ($Mg^{2+}$). Lane 7 consists of DNA treated with 5 μM of Csb2, followed by Proteinase K treatment. A DNA marker is loaded in lane 8, and the respective bands are marked for reference.

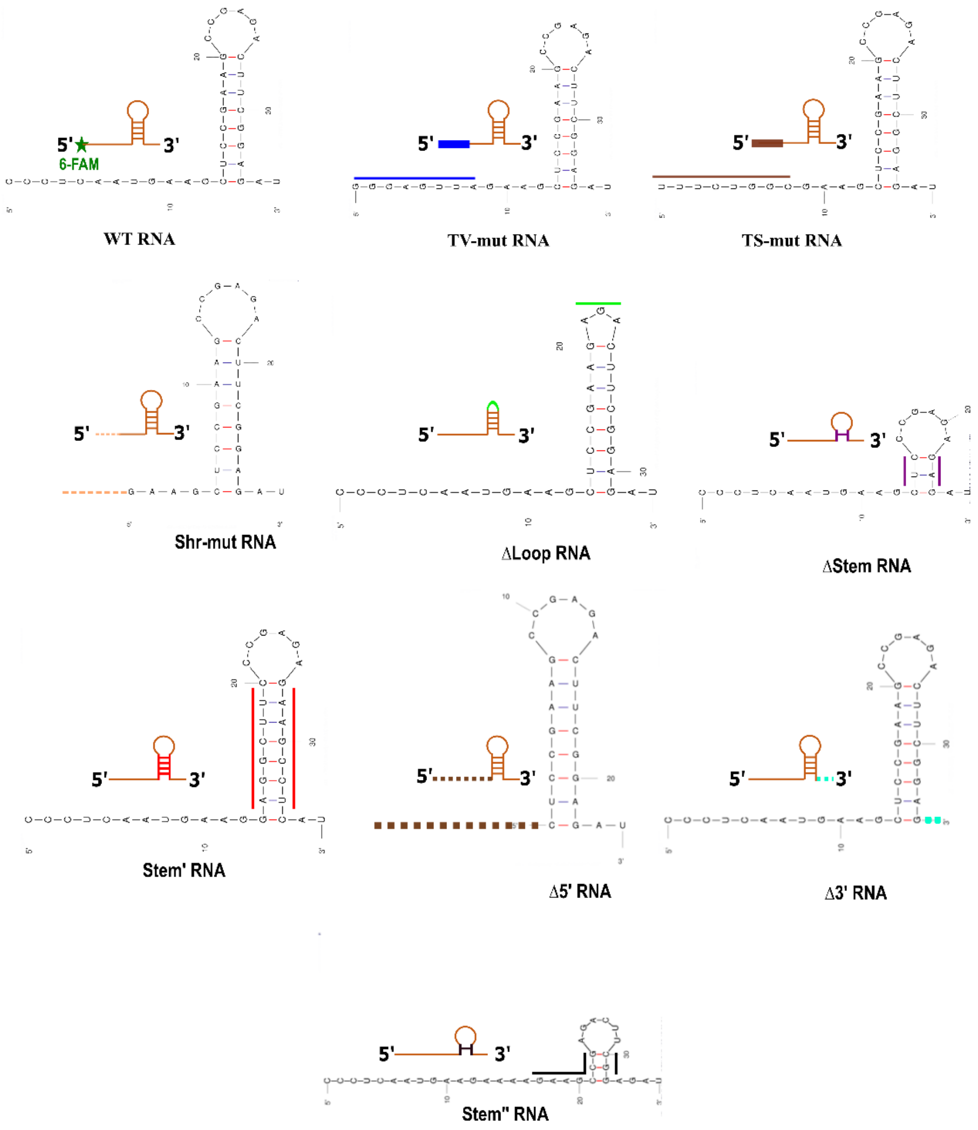


**Figure S2. (A)** Secondary structure representation of the varied repeat RNA substrates that were used to study the influence of sequence and structure of repeat RNA on the nuclease activity of Csb1 and Csb2 (in Figure 2B and S4H). The secondary structure prediction was made using the MFOLD Web Server (1). The respective schematic representations of the RNA used in Figure 2A are indicated along with the predicted secondary structure. TV-mut (blue) and TS-mut (brown) RNA indicate transversion and transition mutations incorporated in the first 8 nt from the 5' end, respectively. Shr-mut RNA (orange dashed lines) indicates a deletion of the first 8 nt from the 5' end. ΔLoop (green) and ΔStem RNA (purple) indicate deletion of the loop and the stem region from the WT substrate, respectively. Stem' RNA (red) represent the variant where the residues of the stem region were interchanged. Δ5' (dark brown dashes) and Δ3' RNA (cyan dashes) indicate deletion of 5' overhang (12 nt from the 5' end) and 3' overhang (2 nt from the 3'end), respectively. Stem" RNA (black) variant indicates destabilisation of the stem region, yet, the total length with respect to the WT remains unaltered (36 nt).

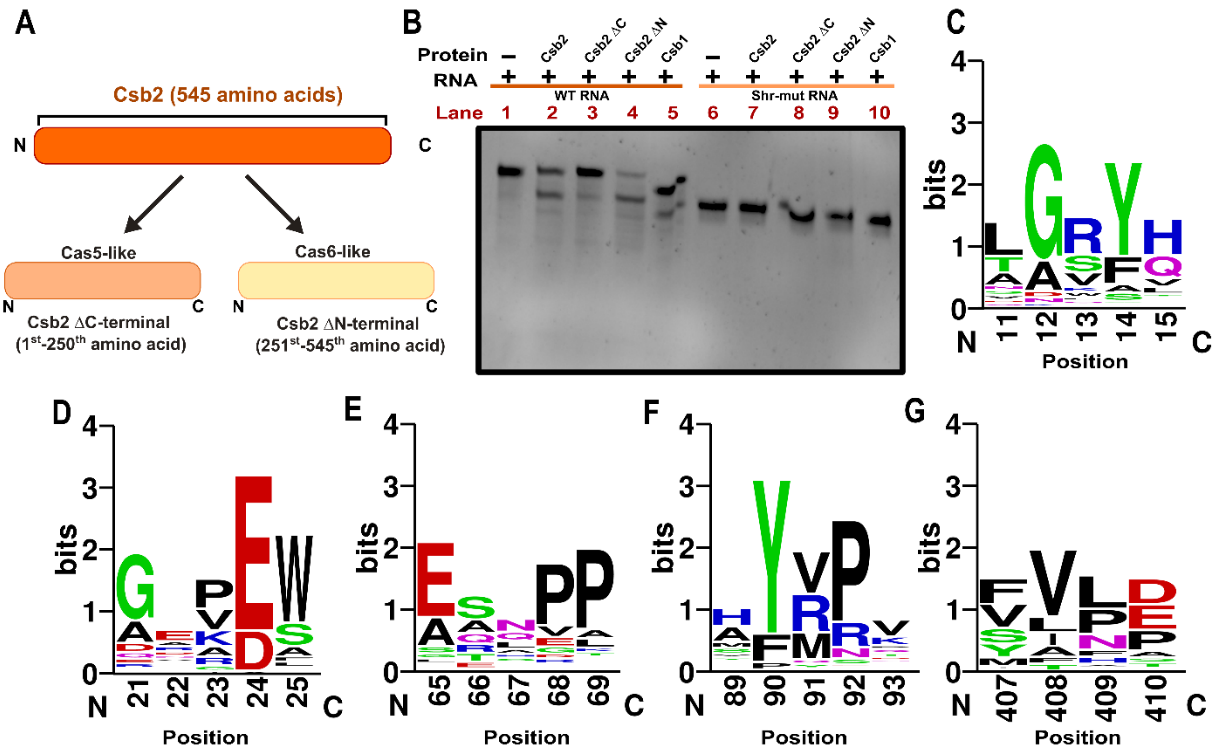


**Figure S3. (A)** A schematic representation of the domain assignment of Csb2 (orange) that shows the presence of Cas5-like and Cas6-like domains. FFAS analysis (2) revealed that the N-terminal region of Csb2 from $1^{st}$ -$250^{th}$ amino acid (peach), namely, Csb2 ΔC-terminal, is homologous to Cas5 whereas the rest from $251^{st}$ – $545^{th}$ amino acid (light yellow), namely, Csb2 ΔN-terminal, is homologous to Cas6. **(B)** An EtBr stained 20% denaturing PAGE probes the RNase activity of 1 µM Csb2 ΔC (lanes 3, 8) and ΔN (lanes 4, 9) domains on 0.5 µM $RNA_{WT}$ and Shr-mut RNA, respectively. As a control, Lanes 2, 7 show the RNase activity of Csb2 and lanes 5,10 show the RNase activity of Csb1 on $RNA_{WT}$ and Shr-mut RNA, respectively. **(C-G)** The sequence logo depicts the conservation of amino acids Y14 (C), E24 (D), E65 (E), Y90 (F), D410 (G) across Csb2 proteins (57 organisms) in the type I-G CRISPR-Cas system. The X-axis indicates the amino acid residue number, whereas, the Y-axis indicates the extent of its conservation (in bits).

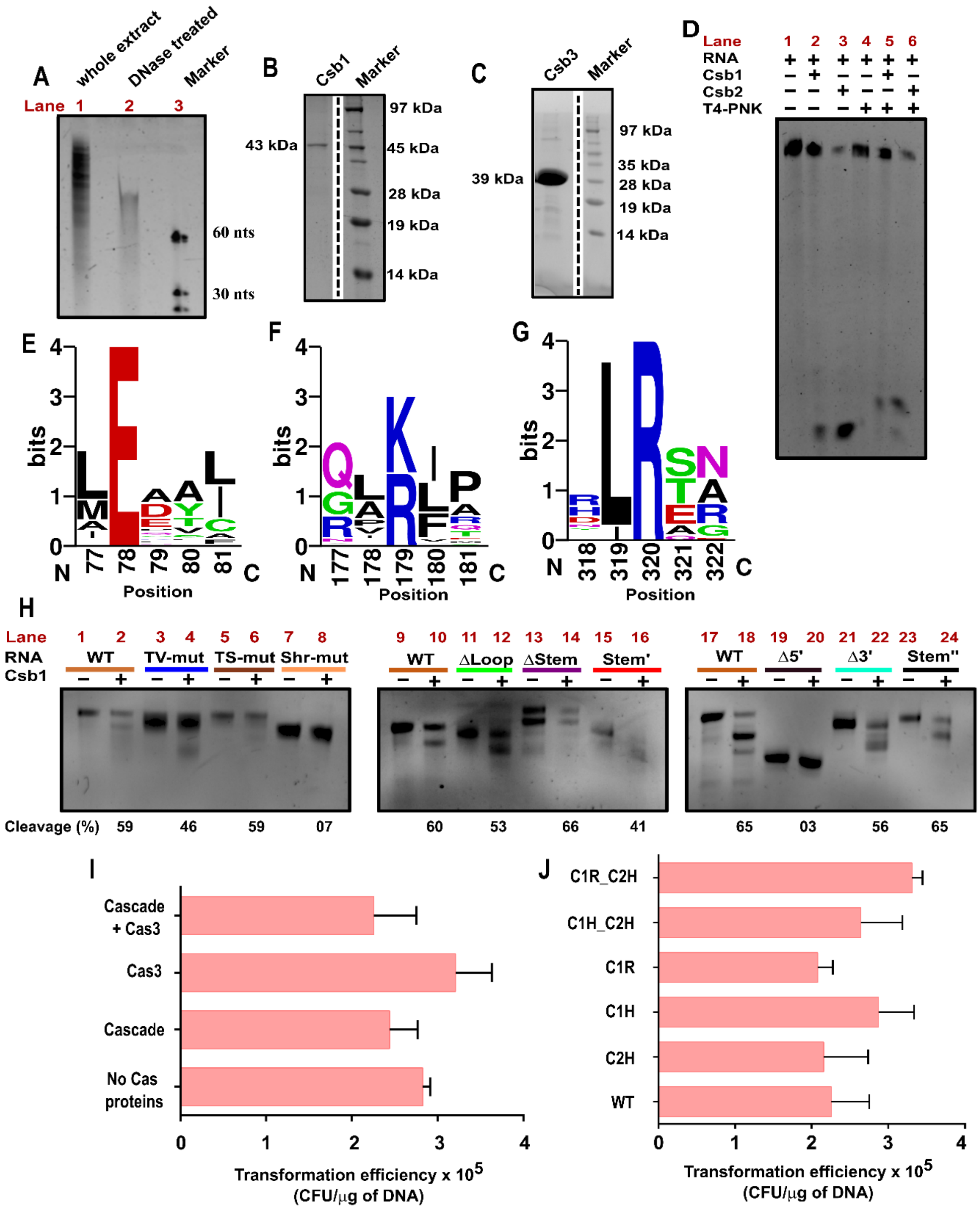
A
whole extract
DNase treated
Marker
Lane 1 2 3
60 nts
30 nts
B
Csb1
Marker
43 kDa
97 kDa
45 kDa
28 kDa
19 kDa
14 kDa
C
Csb3
Marker
39 kDa
97 kDa
35 kDa
28 kDa
19 kDa
14 kDa
D
Lane 1 2 3 4 5 6
RNA
Csb1
Csb2
T4-PNK
E
bits
Position
F
bits
Position
G
bits
Position
H
Lane
RNA
Csb1
WT
TV-mut
TS-mut
Shr-mut
ΔLoop
ΔStem
Stem'
Δ5'
Δ3'
Stem''
Cleavage (%) 59 46 59 07 60 53 66 41 65 03 56 65
I
Cascade + Cas3
Cas3
Cascade
No Cas proteins
Transformation efficiency x 10^5 (CFU/μg of DNA)
J
C1R_C2H
C1H_C2H
C1R
C1H
C2H
WT
Transformation efficiency x 10^5 (CFU/μg of DNA)

**Figure S4. (A)** An EtBr stained 20% denaturing PAGE shows the presence of RNA that was isolated from the purified Cascade complex. Lane 1 indicates the nucleic acid extract from the Cascade complex and lane 2 indicates the same treated with DNase to confirm the presence of RNA. An ssDNA marker is loaded onto lane 3 for reference. **(B, C)** A 15% SDS PAGE shows the purified 43 kDa Csb1 protein (marked as Csb1) and 39 kDa Csb3 protein (marked as Csb3), respectively. A protein marker is loaded on the left to validate the molecular sizes. The vertical broken line indicates discontinuity between the lanes, which is introduced for the sake of clarity. **(D)** A 20% denaturing PAGE is shown for deciphering the nature of cleaved RNA product by Csb1 (lane 5). The cleaved RNA product was treated with T4 PNK at pH 5.2 in the absence of ATP to replace terminal phosphates with hydroxyl groups. The difference in migration between T4 PNK treated and untreated samples is attributed to the charge difference between these groups. Similarly, Csb2 cleaved RNA fragment was also treated with T4 PNK and loaded in lane 6 for comparison. **(E-G)** The sequence logo depicts the conservation of amino acids, namely, E78 (E), R179 (F), and R320 (G) in Csb1 across the type I-G CRISPR-Cas system (57 organisms), respectively. The X-axis indicates the amino acid residue number, whereas, the Y-axis indicates the extent of its conservation (in bits). **(H)** An EtBr stained 20% denaturing PAGE shows the nuclease activity of Csb1 on the variants of CRISPR repeat RNA as described in Figures 2A and S2. **(I)** Transformation efficiency of the strain IG-CR in response to the non-target plasmid is shown. Unlike the target plasmid, the transformation efficiency is not dramatically altered in response to non-target plasmid. Error bar represents standard deviation measured from three independent trials. (**J**) Transformation efficiency of the strain IG-CR in response to the non-target plasmid and in the presence of WT and mutant Cascade complex such as C2H (Csb2 H520A Cascade), C1H (Csb1 H122A Cascade), C1R (Csb1 R123A Cascade), C1H_C2H (Csb1 H122A, Csb2 H520A Cascade), C1R_C2H – (Csb1 R123A, Csb2 H520A Cascade) is shown. The error bars represent standard deviation observed across three independent trials.

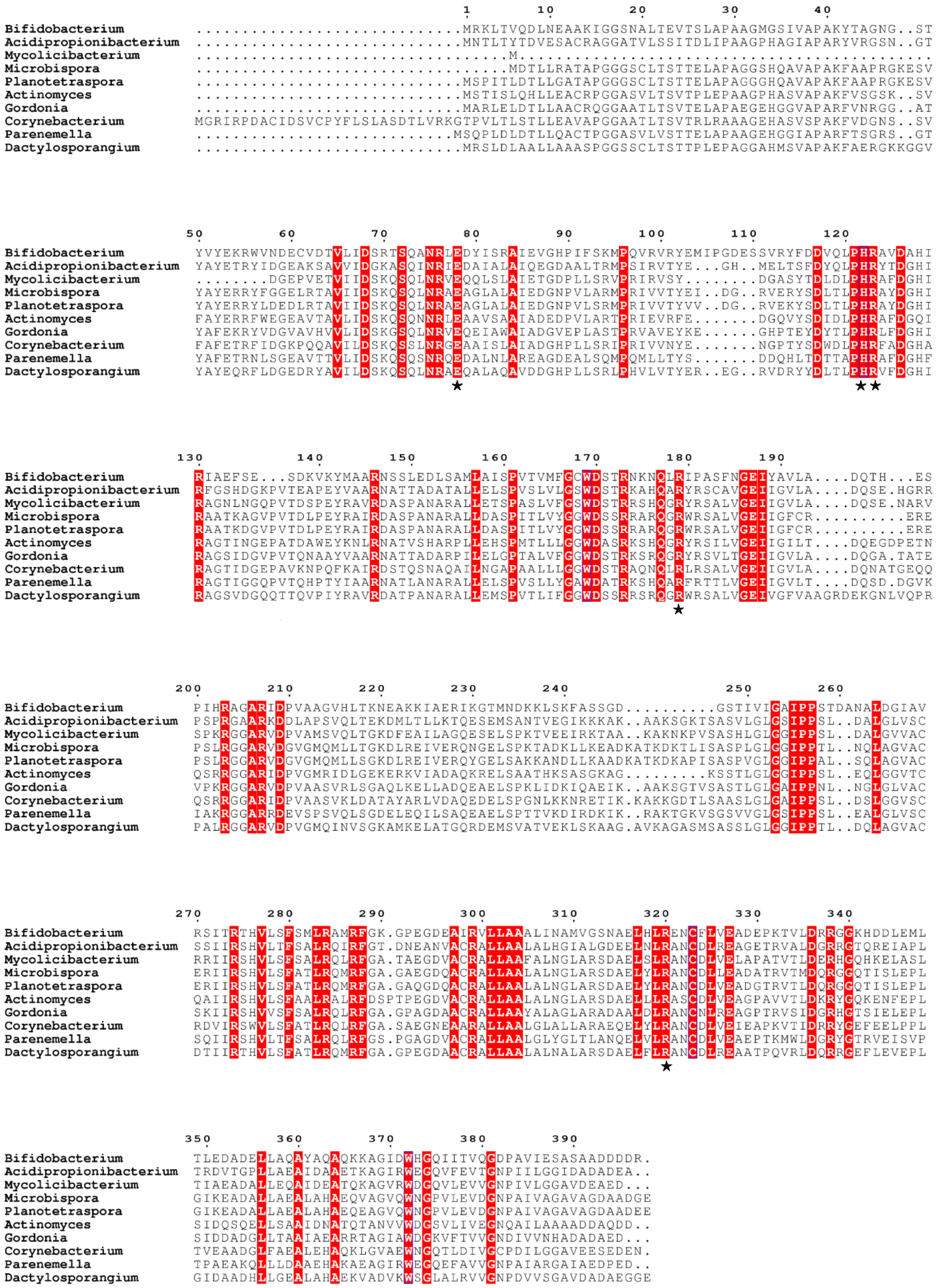
Bifidobacterium ............................MRKLTVQDLNEAAKIGGSNALTEVTSLAPAAGMGSIVAPAKYTAGNG..ST
Acidipropionibacterium ............................MNTLTYTDVESACRAGGATVLSSITDLIPAAGPHAGIAPARYVRGSN..GT
Mycolicibacterium ............................M.......................................................
Microbispora .................................MDTLLRATAPGGGSCLTSTTELAPAGGSHQAVAPAKFAAPRGKESV
Planotetraspora ............................MSPITLDTTLLGATAPGGGSCLTSTTELAPAGGGHQAVAPAKFAAPRGKESV
Actinomyces ............................MSTISLQHLLEACRPGGASVLTSVTPLEPAAGPHASVAPAKFVSGSK..SV
Gordonia ............................MARLELDTLLAACRQGGAATLTSVTELAPAEGEHGGVAPARFVNRGG..AT
Corynebacterium MGRIRPDACIDSVCPYFLSLASDTLVRKGTPVLTLSTLLEAVAPGGAATLTSVTRLRAAAGEHASVSPAKFVDGNS..SV
Parenemella ...........................MSQPLDLDTLLQACTPGGASVLVSTTELAPAAGEHGGIAPARFTSGRS..GT
Dactylosporangium ............................MRSLDLAALLAAASPGGSSCLTSTTPLEPAGGAHMSVAPAKFAERGKKGGV
Bifidobacterium YVYEKRWVNDECVDTVLIDSRTSQANRLEDYISRAIEVGHPIFSKMPQVRVRYEMIPGDESSVRYFDDVQLPHRAVDAHI
Acidipropionibacterium YAYETRYIDGEAKSAVVIDGKASQINRIEDAIALAIQEGDAALTRMPSIRVTYE...GH..MELTSFDYQLPHRYTDGHI
Mycolicibacterium ........DGEPVETVIIDSKQSQLNRVEQQLSLAIETGDPLLSRVPRIRVSY.......DGASYTDLDLPHRAFDGHI
Microbispora YAYERRYFGGELRTAVIIDSKQSQLNRAEAGLALAIEDGNPVLARMPRIVVTYEI..DG..RVERYSDLTLPHRAYDGHI
Planotetraspora YAYERRYLDEDLRTAVIIDSKQSQLNRAEAGLALAIEDGNPVLSRMPRIVVTYVV..DG..RVEKYSDLTLPHRAYDGHI
Actinomyces FAYERRFWEGEAVTAVLIDSKQSQNNRLEAAVSAAIADEDPVLARTPRIEVRFE.......DGQVYSDIDLPHRAFDGQI
Gordonia YAFEKRYVDGVAHVVLIDSKGSQLNRVEQEIAWAIADGVEPLASTPRVAVEYKE......GHPTEYDYTLPHRLFDGHI
Corynebacterium FAFETRFIDGKPQQAVILDSKQSSLNRGEAAISLAIADGHPLLSRIPRIVVNYE.......NGPTYSDWDLPHRFADGHA
Parenemella YAFETRNLSGEAVTTVLIDSKQSQSNRQEDALNLAREAGDEALSQMPQMLLTYS.......DDQHLTDTTAPHRAFDGHF
Dactylosporangium YAYEQRFLDGEDRYAVILDSKQSQLNRAEQALAQAVDDGHPLLSRLPHVLVTYER..EG..RVDRYYDLTLPHRVFDGHI
Bifidobacterium RIAEFSE...SDKVKYMAARNSSLEDLSAMLAISPVTVMFGCWDSTRNKNQLRIPASFNGEIYAVLA....DQTH...ES
Acidipropionibacterium RFGSHDGKPVTEAPEYVAARNATTADATALLELSPVSLVLGSWDSTRKAHQARYRSCAVGEIIGVLA....DQSE.HGRR
Mycolicibacterium RAGNLNGQPVTDSPEYRAVRDASPANARALLETSPASLVFGSWDSTRRSHQGRYRSALVGEIIGVLA....DQSE.NARV
Microbispora RAATKAGVPVTDLPEYRAIRDASPANARALLDASPITLVYGGWDSSRRARQGRWRSALVGEIIGFCR.........ERE
Planotetraspora RAATKDGVPVTDLPEYRAIRDASPANARALLDASPITLVYGGWDSSRRARQGRWRSALVGEIIGFCR.........ERE
Actinomyces RAGTINGEPATDAWEYKNLRNATVSHARPLLEHSPMTLLLGGWDASRKSHQGRYRSILVGEIIGILT....DQEGDPETN
Gordonia RAGSIDGVPVTQNAAYVAARNATTADARPILELGPTALVFGGWDSTRKSRQGRYRSVLTGEIIGVLA....DQGA.TATE
Corynebacterium RAGTIDGEPAVKNPQFKAIRDSTQSNAQAILNGAPAALLLGGWDSTRAQNQLRLRSALVGEIIGVLA....DQNATGEQQ
Parenemella RAGTIGGQPVTQHPTYIAARNATLANARALLELSPVSLLYGAWDATRKSHQARFRTTLVGEIIGVLT....DQSD.DGVK
Dactylosporangium RAGSVDGQQTTQVPIYRAVRDATPANARALLEMSPVTLIFGGWDSSRRSRQGRWRSALVGEIVGFVAAGRDEKGNLVQPR
Bifidobacterium PIHRAGARIDPVAAGVHLTKNEAKKIAERIKGTMNDKKLSKFASSGD..........GSTIVIGAIPPSTDANALDGIAV
Acidipropionibacterium PSPRGAARKDDLAPSVQLTEKDMLTLLKTQESEMSANTVEGIKKKAK..AAKSGKTSASVLGLGSIPPSL..DALGLVSC
Mycolicibacterium SPKRGGARVDPVAMSVQLTGKDFEAILAGQESELSPKTVEEIRKTAA..KAKNKPVSASHLGLGGIPPSL..DALGVVAC
Microbispora PSLRGGARVDGVGMQMLLTGKDLREIVERQNGELSPKTADKLLKEADKATKDKTLISASPLGLGGIPPTL..NQLAGVAC
Planotetraspora PSLRGGARVDGVGMQMLLSGKDLREIVERQYGELSAKKANDLLKAADKATKDKAPISASPVGLGGIPPAL..SQLAGVAC
Actinomyces QSRRGGARIDPVGMRIDLGEKERKVIADAQKRELSAATHKSASGKAG.........KSSTLGLGGIPPSL..EQLGGVTC
Gordonia VPKRGGARVDPVAASVRLSGAQLKELLADQEAELSPKLIDKIQAEIK..AAKSGTVSASTLGLGAIPPNL..NGLGLVAC
Corynebacterium QSRRGGARIDPVAASVKLDATAYARLVDAQEDELSPGNLKKNRETIK.KAKKGDTLSAASLGLGAIPPSL..DSLGGVSC
Parenemella IAKRGGARRDEVSPSVQLSGDELEQILSAQEAELSPTTVKDIRDKIK..RAKTGKVSGSVVGLGSIPPSL..EALGLVSC
Dactylosporangium PALRGGARVDPVGMQINVSGKAMKELATGQRDEMSVATVEKLSKAAG.AVKAGASMSASSLGLGGIPPTL..DQLAGVAC
Bifidobacterium RSITRTHVLSFSMLRAMRFGK.GPEGDEAIRVLLAAALINAMVGSNAELHLRENCFLVEADEPKTVLDRRGGKHDDLEML
Acidipropionibacterium SSIIRSHVLTFSALRQIRFGT.DNEANVACRALLAAALALHGIALGDEELNLRANCDLREAGETRVALDGRRGTQREIAPL
Mycolicibacterium RRIIRSHVLSFSALRQLRFGA.TAEGDVACRALLAAFALNGLARSDAELSLRANCDLVELAPATVTLDERHGQHKELASL
Microbispora ERIIRSHVLSFATLRQMRFGA.GAEGDQACRALLAALALNGLARSDAELYLRANCDLLEADATRVTMDQRGGQTISLEPL
Planotetraspora ERIIRSHVLSFATLRQMRFGA.GAQGDQACRALLAALALNGLARSDAELYLRANCDLVEADGTRVTLDQRGGQTISLEPL
Actinomyces QAIIRSHVLSFAALRALRFDSPTPEGDVACRALLAALALNGLARSDAELLLRASCDLVEAGPAVVTLDKRYGQKENFEPL
Gordonia SKIIRSHVVSFSALRQLRFGA.GPAGDAACRALLAAYALAGLARADAALDLRANCNLREAGPTRVSIDGRHGTSIELEPL
Corynebacterium RDVIRSWVLSFATLRQLRFGA.SAEGNEAARALLAALGLALLARAEQELYLRANCDLVEIEAPKVTIDRRYGEFEELPPL
Parenemella SQIIRSHVLTFSALRQLRFGS.PGAGDVACRALLAALGLYGLTLANQELVLRANCDLVEAEPTKMWLDGRYGTRVEISVP
Dactylosporangium DTIIRTHVLSFATLRQMRFGA.GPEGDAACRALLAALALNALARSDAELFLRANCDLREAATPQVRLDQRRGEFLEVEPL
Bifidobacterium TLEDADELLAQAYAQAQKKAGIDWHGQIITVQGDPAVIESASAADDDDR.
Acidipropionibacterium TRDVTGPLLAEAIDAAETKAGIRWEGQVFEVTGNPIILGGIDADADEA..
Mycolicibacterium TIAEADALLEQAIDEATQKAGVRWDGQVLEVVGNPIVLGGAVDEAED...
Microbispora GIKEADALLAEALAHAEQVAGVQWNGPVLEVDGNPAIVAGAVAGDAADGE
Planotetraspora GIKEADALLAEALAHAEQEAGVQWNGPVLEVDGNPAIVAGAVAGDAADEE
Actinomyces SIDQSQELLSAAIDNATQTANVVWDGSVLIVEGNQAILAAAADDAQDD..
Gordonia SIDDADGLLTAAIAEARRTAGIAWDGKVFTVVGNDIVVNHADADAED...
Corynebacterium TVEAADGLFAEALEHAQKLGVAEWNGQTLDIVGCPDILGGAVEESEDEN.
Parenemella TPAEAKQLLLDAAEHAKAEAGIRWEGQEFAVVGNPAIARGAIAEDPED..
Dactylosporangium GIDAADHLLGEALAHAEKVADVKWSGLALRVVGNPDVVSGAVDADAEGGE

**Figure S5.** Multiple sequence alignment corresponding to Csb1/I-G from select organisms is shown. The sequences were obtained from *Bifidobacterium animalis*, *Acidipropionibacterium virtanenii*, *Mycolicibacterium hassiacum* DSM 44199, *Microbispora* sp. CL1-1, *Planotetraspora phitsanulokensis*, *Actinomyces* sp. ZJ308, *Gordonia iterans*, *Corynebacterium flavescens*, *Parenemella sanctibonifatiensis*, and *Dactylosporangium aurantiacum* and were aligned using MUSCLE (3). Amino acid residues that are strictly conserved are highlighted in red and the conserved residues used for the site directed mutagenesis studies (E78, H122, R123, R179 and R320) are indicated using a star symbol. The alignment is highlighted based on the extent of conservation using ESpript (4)

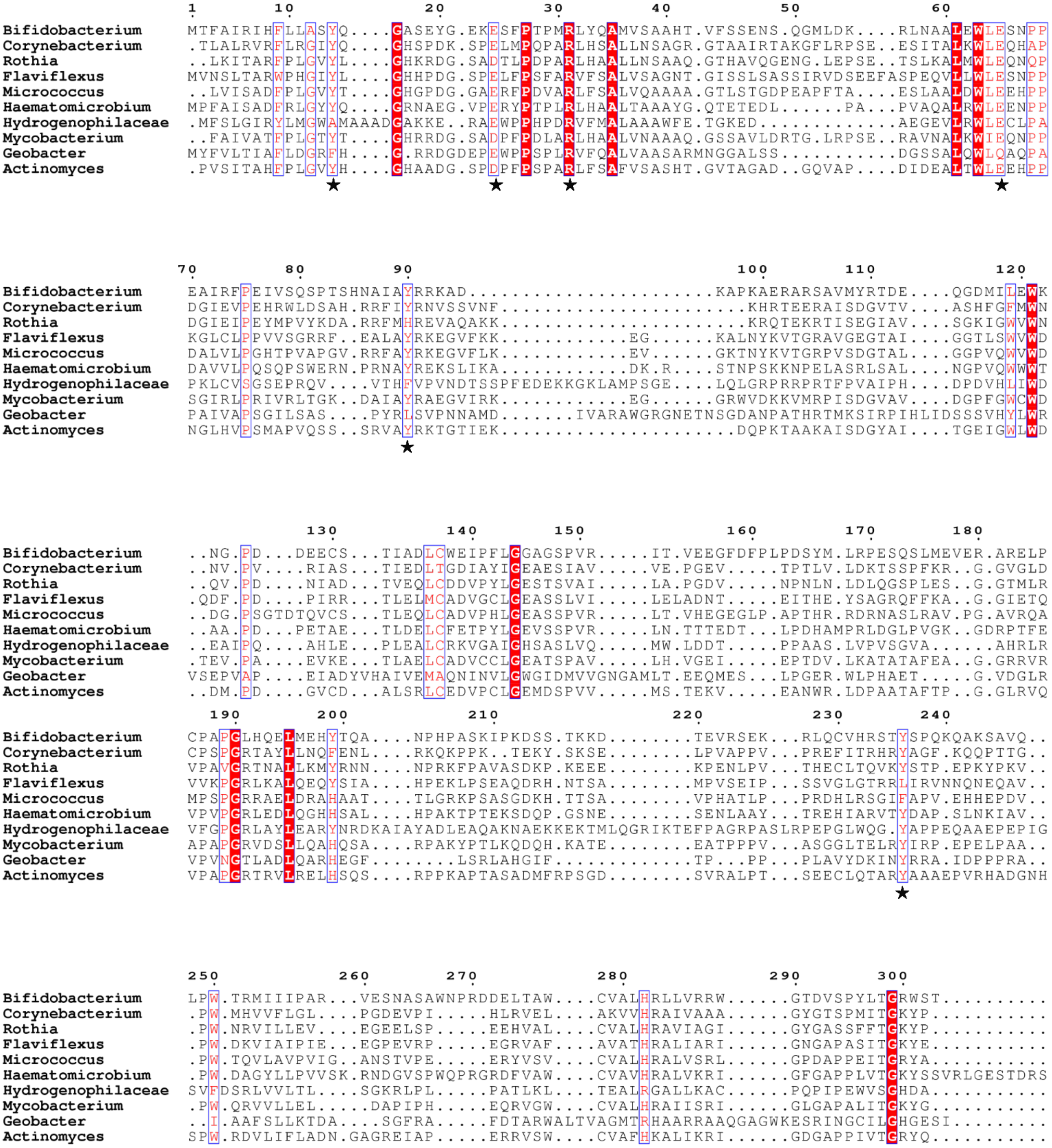
1 10 20 30 40 50 60
Bifidobacterium MTFAIRIHFLLASYQ....GASEYG.EKESFPTPMRLYQAMVSAAHT.VFSSENS.QGMLDK....RLNAALEWLESNPP
Corynebacterium .TLALRVRFLRGIYQ....GHSPDK.SPELMPQPARLHSALLNSAGR.GTAAIRTAKGFLRPSE..ESITALKWLEQHAP
Rothia ..LKITARFPLGVYL....GHKRDG.SADTLPDPARLHAALLNSAAQ.GTHAVQGENG.LEPSE..TSLKALMWLEQNQP
Flaviflexus MVNSLTARWPHGIYL....GHHPDG.SPELFPSFARVFSALVSAGNT.GISSLSASSIRVDSEEFASPEQVLLWLESNPP
Micrococcus ..LVISADFPLGVYT....GHGPDG.GAERFPDVARLFSALVQAAAA.GTLSTGDPEAPFTA....ESLAALDWLEEHPP
Haematomicrobium MPFAISADFRLGYYQ....GRNAEG.VPERYPTPLRLHAALTAAAYG.QTETEDL......PA....PVAQALRWLEENPP
Hydrogenophilaceae .MFSLGIRYLMGWAMAAADGAKKE..RAEWPPHPDRVFMALAAAWFE.TGKED............AEGEVLRWLECLPA
Mycobacterium ..FAIVATFPLGTYT....GHRRDG.SADPFPDLARLHAALVNAAAQ.GSSAVLDRTG.LRPSE..RAVNALKWIEQNPP
Geobacter MYFVLTIAFLDGRFH....G.RRDGDEPEWPPSPLRVFQALVAASARMNGGALSS..........DGSSALQWLQAQPA
Actinomyces .PVSITAHFPLGVYH....GHAADG.SPDPFPSPARLFSAFVSASHT.GVTAGAD..GQVAP....DIDEALTWLEEHPP
70 80 90 100 110 120
Bifidobacterium EAIRFPEIVSQSPTSHNAIAYRRKAD......................KAPKAERARSAVMYRTDE....QGDMILEWK
Corynebacterium DGIEVPEHRWLDSAH.RRFIYRNVSSVNF......................KHRTEERAISDGVTV....ASHFGFMWN
Rothia DGIEIPEYMPVYKDA.RRFMHREVAQAKK......................KRQTEKRTISEGIAV....SGKIGWVWN
Flaviflexus KGLCLPPVVSGRRF..EALAYRKEGVFKK............EG.....KALNYKVTGRAVGEGTAI....GGTLSWVWD
Micrococcus DALVLPGHTPVAPGV.RRFAYRKEGVFLK............EV.....GKTNYKVTGRPVSDGTAL....GGPVQWVWD
Haematomicrobium DAVVLPQSQPSWERN.PRNAYREKSLIKA............DK.R...STNPSKKNPELASRLSAL....NGPVQWWWT
Hydrogenophilaceae PKLCVSGSEPRQV....VTHFVPVNDTSSPFEDEKKGKLAMPSGE...LQLGRPRRPRTFPVAIPH....DPDVHLIWD
Mycobacterium SGIRLPRIVRLTGK..DAIAYRAEGVIRK............EG.....GRWVDKKVMRPISDGVAV....DGPFGWCWD
Geobacter PAIVAPSGILSAS....PYRLSVPNNAMD......IVARAWGRGNETNSGDANPATHRTMKSIRPIHLIDSSSVHYLWR
Actinomyces NGLHVPSMAPVQSS..SRVAYRKTGTIEK.....................DQPKTAAKAISDGYAI....TGEIGWLWD
130 140 150 160 170 180
Bifidobacterium ..NG.PD...DEECS...TIADLCWEIPFLGGAGSPVR.....IT.VEEGFDFPLPDSYM.LRPESQSLMEVER.ARELP
Corynebacterium ..NV.PV....RIAS...TIEDLTGDIAYIGEAESIAV....VE.PGEV.....TPTLV.LDKTSSPFKR..G.GVGLD
Rothia ..QV.PD....NIAD...TVEQLCDDVPYLGESTSVAI....LA.PGDV.....NPNLN.LDLQGSPLES..G.GTMLR
Flaviflexus .QDF.PD....PIRR...TLELMCADVGCLGEASSLVI....LELADNT.....EITHE.YSAGRQFFKA..G.GIETQ
Micrococcus ..DG.PSGTDTQVCS...TLEQLCADVPHLGEASSPVR....LT.VHEGEGLP.APTHR.RDRNASLRAV.PG.AVRQA
Haematomicrobium ..AA.PD...PETAE...TLDELCFETPYLGEVSSPVR....LN.TTTEDT...LPDHAMPRLDGLPVGK..GDRPTFE
Hydrogenophilaceae ..EAIPQ....AHLE...PLEALCRKVGAIGHSASLVQ....MW.LDDT.....PPAAS.LVPVSGVA......AHRLR
Mycobacterium .TEV.PA....EVKE...TLAELCADVCCLGEATSPAV....LH.VGEI.....EPTDV.LKATATAFEA..G.GRRVR
Geobacter VSEPVAP....EIADYVHAIVEMAQNINVLGWGIDMVVGNGAMLT.EEQMES...LPGER.WLPHAET.....G.VDGLR
Actinomyces ..DM.PD....GVCD...ALSRLCEDVPCLGEMDSPVV....MS.TEKV.....EANWR.LDPAATAFTP..G.GLRVQ
190 200 210 220 230 240
Bifidobacterium CPAPGLHQELMEHYTQA....NPHPASKIPKDSS.TKKD........TEVRSEK...RLQCVHRSTYSPQKQAKSAVQ..
Corynebacterium CPSPGRTAYLLNQFENL....RKQKPPK..TEKY.SKSE........LPVAPPV...PREFITRHRYAGF.KQQPTTG..
Rothia VPAVGRTNALLKMYRNN....NPRKFPAVASDKP.KEEE........KPENLPV...THECLTQVKYSTP.EPKYPKV..
Flaviflexus VVKPGRLKALQEQYSIA....HPEKLPSEAQDRH.NTSA........MPVSEIP...SSVGLGTRRLIRVNNQNEQAV..
Micrococcus MPSPGRRAELDRAHAAT....TLGRKPSASGDKH.TTSA........VPHATLP...PRDHLRSGIFAPV.EHHEPDV..
Haematomicrobium VPVPGRLEDLQGHHSAL....HPAKTPTEKSDQP.GSNE........SENLAAY...TREHIARVTYDAP.SLNKIAV..
Hydrogenophilaceae VFGPGRLAYLEARYNRDKAIAYADLEAQAKNAEKKEKTMLQGRIKTEFPAGRPASLRPEPGLWQG.YAPPEQAAEPEPIG
Mycobacterium APAPGRVDSLLQAHQSA....RPAKYPTLKQDQH.KATE........EATPPPV...ASGGLTELRYIRP.EPELPAA..
Geobacter VPVNGTLADLQARHEGF........LSRLAHGIF.............TP...PP...PLAVYDKINYRRA.IDPPPRA..
Actinomyces VPAPGRTRVLRELHSQS....RPPKAPTASADMFRPSGD........SVRALPT...SEECLQTARYAAAEPVRHADGNH
250 260 270 280 290 300
Bifidobacterium LPW.TRMIIIPAR...VESNASAWNPRDDELTAW....CVALHRLLVRRW......GTDVSPYLTGRWST..........
Corynebacterium .PW.MHVVFLGL....PGDEVPI.....HLRVEL....AKVVHRAIVAAA......GYGTSPMITGKYP...........
Rothia .PW.NRVILLEV....EGEELSP.....EEHVAL....CVALHRAVIAGI......GYGASSFFTGKYP...........
Flaviflexus .PW.DKVIAIPIE...EGPEVRP.....EGRVAF....AVATHRALIARI......GNGAPASITGKYE...........
Micrococcus .PW.TQVLAVPVIG..ANSTVPE.....ERYVSV....CVALHRALVSRL......GPDAPPEITGRYA...........
Haematomicrobium .PW.DAGYLLPVVSK.RNDGVSPWQPRGRDFVAW....CVAVHRALVKRI......GFGAPPLVTGKYSSVRLGESTDRS
Hydrogenophilaceae SVFDSRLVVLTL....SGKRLPL.....PATLKL....TEALRGALLKAC......PQPIPEWVSGHDA...........
Mycobacterium .PW.QRVVLLEL.....DAPIPH.....EQRVGW....CVALHRAIISRI......GLGAPALITGKYG...........
Geobacter ..I.AAFSLLKTDA....SGFRA.....FDTARWALTVAGMTRHAARRAAQGAGWKESRINGCILGHGESI.........
Actinomyces SPW.RDVLIFLADN.GAGREIAP.....ERRVSW....CVAFHKALIKRI......GDGAPPIVTGRYQ...........

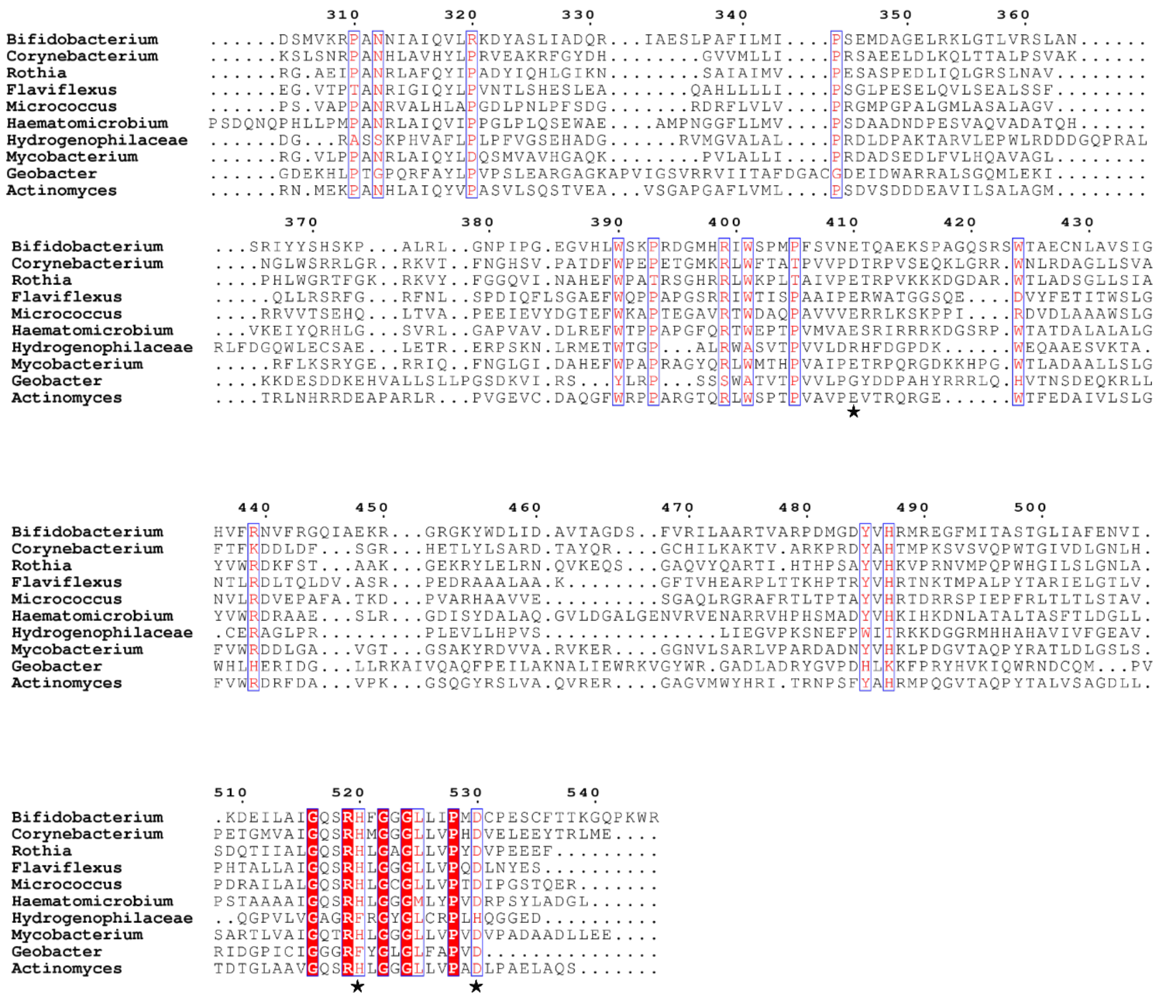


**Figure S6.** Multiple sequence alignment corresponding to Csb2/I-G from select organisms is shown. The sequences were obtained from *Bifidobacterium animalis*, *Corynebacterium* sp. HMSC063G05, *Rothia* sp. HMSC036D11, *Flaviflexus massiliensis*, *Micrococcus* sp. KBS0714, *Haematomicrobium sanguinis*, *Hydrogenophilaceae bacterium* CG1_02_62_390, *Mycobacterium hassiacum*, *Geobacter sulfurreducens*, and *Actinomyces oris* and were aligned using MUSCLE (3). Amino acid residues that are completely conserved are highlighted in red, those conserved/similar across the groups are highlighted in blue box and red text and the conserved residues used for the site directed mutagenesis studies (Y14, E24, R31, E65, Y90, Y236, E410, H520, and D530) are indicated using a star symbol. The alignment is highlighted based on the extent of conservation using ESpript (4)

#### Supplementary Tables

Table S1: **Sequences of the various oligonucleotides used in the study**

| Oligo name | Sequence (5’-3’) | Description |
|---|---|---|
| Csb1-13S-R FP | AACCTGTACTTCCAATCCAATGCAATGATGCGCAAACTCACAGTACAAG | Amplification of gene encoding Csb1 from *B. animalis* with restriction sites SspI for 13S-R plasmid. |
| Csb1-13S-R RP | TTATCCACTTCCAATGTTATTATTATCATCTATCATCGTCGTCAGCAGCG | |
| Csb2-1R FP | TACTTCCAATCCAATGCAATGACGTTCGCGATTCGTATCCAC | Amplification of gene encoding Csb2 from *B. animalis* with restriction sites SspI for 1R plasmid. |
| Csb2-1R RP | TTATCCACTTCCAATGTTATTATCATCTCCATTTCGGTTGCCCTTT | |
| Csb3-pQE2 FP | ACATCACCATCACCATCACCATATG ATGAGCGTTCTGCGAATTCC | Amplification of gene encoding Csb3 from *B. animalis* with restriction sites NdeI and HindIII for pQE2 plasmid |
| Csb3-pQE2 RP | AGTCCAAGCTCAGCTAATTAAGCTT CTAGAGCCGTATGAGCTGCC | |
| Cascade-pQE2 FP | ACATCACCATCACCATCACCATATGATGACGTTCGCGATTCGTATCCAC | Amplification of gene encoding Csb2 to assemble the Cascade operon from *B. animalis* with restriction sites NdeI and HindIII for pQE2 plasmid |
| Csb2-Csb1 overlap RP | ACTGTGAGTTTGCGCATCATGGTATATCTCCTTCTTAAATTATCATCTCCATTT CGGTT | Amplification of gene encoding Csb2 and overlapping region of Csb1, to assemble the Cascade operon from *B. animalis* in pQE2 vector |
| Csb2-1 overlap FP | AACCGAAATGGAGATGATAATTTAAGAAGGAGATATACCATGATGCGCAAA CTCACAGT | Amplification of gene encoding Csb1 and overlapping region of Csb2, to assemble the Cascade operon from *B. animalis* in pQE2 vector |
| Csb1-Csb3 overlap RP | GGAATTCGCAGAACGCTCATGGTATATCTCCTTCTTAAATTATCATCTATCAT CGTCG | Amplification of gene encoding Csb1 and overlapping region of Csb3, to assemble the Cascade operon from *B. animalis* in pQE2 vector |
| Csb1-Csb3 overlap FP | ACGACGATGATAGATGATAATTTAAGAAGGAGATATACCATGAGCGTTCTGC GAATTCC | Amplification of gene encoding Csb3 and overlapping region of Csb1, to assemble the Cascade operon from *B. animalis* in pQE2 vector |

| | | |
|---|---|---|
| Csb2 ΔN-terminal-1R FP | TACTTCCAATCCAATGCAACTCGTATGATTATCATTCCCGCGC | Amplification of gene encoding Csb2 ΔN-terminal from *B. animalis* with restriction sites SspI for 1R plasmid |
| Csb2 ΔN-terminal-1R RP | TTATCCACTTCCAATGTTATTATCACCCTTTCGTTGTGAAACAGCTTTCG | |
| Csb2 ΔC-terminal-1R FP | TACTTCCAATCCAATGCAATGACGTTCGCGATTCGTATCC | Amplification of gene encoding Csb2 ΔC-terminal from *B. animalis* with restriction sites SspI for 1R plasmid |
| Csb2 ΔC-terminal-1R RP | TTATCCACTTCCAATGTTATTATCAAGTCCATGGCAGCTGTACTGC | |
| Csb1 E78A-13S-R RP | TCGGCTGATGTAGTCTGCCAAACGATTGGCCTG | Amplification of gene encoding various point mutants of Csb1 from *B. animalis* with restriction sites SspI for 13S-R plasmid. |
| Csb1 H122A-13S-R RP | GTCCACCGCACGAGCTGGCAATTGAACATCATCGAAATAC | |
| Csb1 R123A-13S-R RP | GCGTCCACCGCAGCATGTGGCAATTGAACATCATCG | |
| Csb1 R179A-13S-R RP | TTGAAGCTGGCGGGAATAGCCAGTTGGTTCTTGTTTCG | |
| Csb1 R320A-13S-R FP | AAGCAATGCGGAATTGCATCTAGCGGAAAACTGCTTCCTTG | |
| Csb2 Y14A-1R RP | ATATTCGCTTGCTCCTTGAGCGGAAGCAAGAAGGAAGTGG | Amplification of gene encoding various point mutants of Csb2 from *B. animalis* with restriction sites SspI for 1R plasmid. |
| Csb2 E24A-1R RP | TGGGAGTTGGAAAAGATGCCTTCTCCCCATATTCG | |
| Csb2 R31A-1R RP | GATACCATTGCCTGATATAAAGCCATGGGAGTTGGAAAAGATTC | |
| Csb2 E65A-1R RP | TCCGGCGGATTCGATGCCAACCATTCGAGTG | |
| Csb2 Y90A-1R RP | CTTTGTCTGCTTTACGCCTAGCAGCAATGGCATTATGGGAGG | |
| Csb2 Y236A-1R RP | GCTTGTTTCTGAGGGCTGGCGGTACTGCGATGTACACA | |
| Csb2 E410A-1R FP | TATGCCATTTTCGGTCAATGCAACACAAGCTGAAAAAAGCC | |
| Csb2 H520A-1R FP | CGGGCAGAGCAGGGCCTTTGGCGGGGGC | |
| Csb2 D530A-1R FP | GCTTACTCATTCCTATGGCTTGTCCCGAAAGCTGTT | |
| Array_CI-pOSIP-CT FP | GAATTCGAGCTCGGTACCCGGGGGATCCTAATACGACTCACTATAGGGGTCACCTTGGATTTCTAACCATGCC | For integration of the CRISPR array from *B. animalis* using clonetegration into P21 *attB* site of IYB5101. |
| Array_CI-pOSIP-CT RP | GCGCCATGCATCTCGAGGCATGCCTGCAGCAGCATAAAGCGGGGAATCTCTCGC | |
| TTT_Target-13S-R FP | GTGTGAAGCTTGCATGCCTGCAGGTCGACTCTAGATTTGACAGTGCGAACACAGTGCAGT | Amplification of TTT PAM containing Target sequence from *B. animalis* with restriction sites PstI and KpnI for 13S-R plasmid. |
| TTT_Target-13S-R RP | CACACGAATTCGAGCTCGGTACCCGGCGCGATCGTCACCGACTGCACTGTGTTCGCACTG | |

| | | |
|---|---|---|
| Cas3-1R FP | TACTTCCAATCCAATGCAATGGAGATGAATGCAACAACCCCAA | Amplification of gene encoding Cas3 from *B. animalis* with restriction sites SspI for 1R plasmid. |
| Cas3-1R RP | TTATCCACTTCCAATGTTATTATCATCGTCCTTCCATGGATATCGTC | |
| T7 Promoter FP | GAAATTAATACGACTCACTATAGG | For annealing the T7 promoter to the DNA constructs of various mutant RNA substrates to be synthesised via *in vitro* RNA transcription |
| TV-Mut RNA RP | ATCTCCGAAGTCTCGGCTTCGGAGCTTCTAACTCCCCCTATAGTGAGTCGTATTAATTTC | For annealing the DNA template to the T7 promoter region for synthesis of mutant RNA substrates via *in vitro* RNA transcription (RNA constructs are described in table S2) |
| TS-Mut RNA RP | ATCTCCGAAGTCTCGGCTTCGGAGCTTCGCCAGAAACCTATAGTGAGTCGTATTAATTTC | |
| Shr-Mut RNA RP | ATCTCCGAAGTCTCGGCTTCGGAGCTTCCCTATAGTGAGTCGTATTAATTTC | |
| ΔLoop RNA RP | ATCTCCGAAGTCTCTTCGGAGCTTCATTGAGGGCCTATAGTGAGTCGTATTAATTTC | |
| ΔStem RNA RP | ATCTCTCTCGGGAGCTTCATTGAGGGCCTATAGTGAGTCGTATTAATTTC | |
| Stem' RNA RP | ATGAGGCTTCTCTCGGGAAGCCTCCTTCATTGAGGGCCTATAGTGAGTCGTATTAATTTC | |
| Δ5' RNA RP | ATCTCCGAAGTCTCGGCTTCGGAGCCTATAGTGAGTCGTATTAATTTC | |
| Δ3' RNA RP | CTCCGAAGTCTCGGCTTCGGAGCTTCATTGAGGGCCTATAGTGAGTCGTATTAATTTC | |
| Stem'' RNA RP | ATCTCCGAAGTCTCGGCTTCTTTTCTTCATTGAGGGCCTATAGTGAGTCGTATTAATTTC | |
| CR_Array | GCTCTTCCCCTGTAGATTAATTAAGCGGCCGCTAATACGACTCACTATAGGGCCCTCAATGAAGCTCCGAAGCCGAGACTTCGGAGATGACAGTGCGAACACAGTGCAGTCGGTGACGATCGCGCCCTCAATGAAGCTCCGAAGCCGAGACTTCGGAGATGACAGTGCGAACACAGTGCAGTCGGTGACGATCGCGCCCTCAATGAAGCTCCGAAGCCGAGACTTCGGAGATGACAGTGCGAACACAGTGCAGTCGGTGACGATCGCGCCCTCAATGAAGCTCCGAAGCCGAGACTTCGGAGATGACAGTGCGAACACAGTGCAGTCGGTGACGATCGCGCCCTCAATGAAGCTCCGAAGCCGAGACTTCGGAGATGACAGTGCGAACACAGTGCAGTCGGTGACGATCGCGCCCTCAATGAAGCTCCGAAGCCGAGACTTCGGAGATCCGCTGAGCAATAACTAGCATAACCCCTTGGGGCCTCTAAACGGGTCTTGAGGGGTTTTTTGGGTACCACGCGTGCGCGCTGATCCGGAAGAGC | DNA sequence encoding type I-G CRISPR array harbouring 7 repeat units interspersed by 6 identical spacer units from *B. animalis* flanking with T7 promoter and T7 terminator, followed by 13S-R vector backbone and BspQ I recognition sites. |

Table S2: **Sequences of the various RNA constructs used in the study** (refer to Figure S2 for predicted secondary structures)

| Name | Sequence (5'-3') | Description |
|---|---|---|
| WT RNA | CCCUCAAUGAAGCUCCGAAGCCGAGACUUCGGAGAU | 5' 6-FAM labelled CRISPR repeat RNA from *B. animalis* |
| TV-Mut RNA | **GGGAGUUA**GAAGCUCCGAAGCCGAGACUUCGGAGAU | Mutant RNA construct where transversion mutations have been introduced in the 8 nt from the 5'end (in bold) |
| TS-Mut RNA | **UUUCUGGC**GAAGCUCCGAAGCCGAGACUUCGGAGAU | Mutant RNA construct where transition mutations have been introduced in the 8 nt from the 5'end (in bold) |
| Shr-Mut RNA | GAAGCUCCGAAGCCGAGACUUCGGAGAU | Mutant RNA construct where 8 nt from the 5'end are deleted. |
| ΔLoop RNA | CCCUCAAUGAAGCUCCGAAGAGACUUCGGAGAU | Mutant RNA construct where the loop region from $21^{st}$ to $23^{rd}$ nt from the 5'end are deleted. |
| ΔStem RNA | CCCUCAAUGAAGCUCCCGAGAGAGAU | Mutant RNA construct where the stem region from 16th to $20^{th}$ and $27^{th}$ to 31st nt from the 5'end are deleted. |
| Stem' RNA | CCCUCAAUGAAGGAGGCUUCCCGAGAGAAGCCUCAU | Mutant RNA construct where the stem region from $13^{th}$ to $20^{th}$ have been interchanged with the residues $27^{th}$ to $34^{th}$ nt, and vice versa |
| Δ5' RNA | CUCCGAAGCCGAGACUUCGGAGAU | Mutant RNA construct where 12 nt from the 5'end are deleted |
| Δ3' RNA | CCCUCAAUGAAGCUCCGAAGCCGAGACUUCGGAG | Mutant RNA construct where 2 nt from the 3'end are deleted |
| Stem'' RNA | CCCUCAAUGAAGAAAAGAAGCCGAGACUUCGGAGAU | Mutant RNA construct where the stem region is deleted keeping the overall length of the repeat RNA intact. |

Table S3: **List of strains used in this study**

| Strain | Genotype | Source |
|---|---|---|
| *E. coli* IYB5101 | F-Δ(*araD-araB*)567 Δ*lacZ4787* (::rrnB-3) λ- rph-1 Δ(*rhaD-rhaB*)568 *hsdR514 araB*::T7-RNAp-*tetA*, TetR | Kind gift from Prof. Udi Qimron |
| *E. coli* BL21(DE3) | F- *ompT hsdsB*(rB-, mB-) *gal dcm* λ(DE3) | NEB |
| *E. coli* TOP10 | F- *mcrA* Δ(*mrr-hsdRMS-mcrBC*) ϕ80*lac*ZΔM15 Δ *lac*X74 *recA1 araD139* Δ(*araleu*)7697 *galU galK rpsL endA1 nupG*, $Str^R$ | Invitrogen |
| *E. coli* IG-CR | IYB5101 P21:: *I-G array*, $Cam^R$ | This study |

Table S4: **List of plasmids used in this study**

| Plasmid name | Description | Source |
|---|---|---|
| pOSIP-CT | *ori* Rγ, *ori* pUC, $Cam^R$, *attP* P21, *ccdB*, λ (*cI857*) encodes P21 integrase under the control of λ promoter (λ pR). | Addgene #45981 |
| pUC19 | *ori* PBR322, $Amp^R$, for gene insertion under the control of lac promoter. | NEB |
| pQE2 | *ori* ColE1, $Amp^R$, expresses gene of interest to synthesize N-terminal 6xHis tagged protein under the IPTG inducible T5 promoter. | |
| pET StrepII TEV LIC cloning vector (p1R) | *ori* pMB1, $Kan^R$, *lacI*, expresses the gene of interest to synthesize N-terminal StrepII tagged protein under the control IPTG inducible promoter (PT7lac). | Addgene #29664 (Scott Gradia) |
| pET StrepII TEV co-transformation cloning vector (p13SR) / pNT | *ori* CloDF13, $Spc^R$, *lacI*, expresses the gene of interest to synthesize N-terminal StrepII tagged protein under the control IPTG inducible promoter (PT7lac). | Addgene #48328 (Scott Gradia) |
| pCsb2/I-G | *csb2* gene from *B. animalis* encoding N-terminal Strep tagged protein inserted in p1R plasmid. Kanamycin resistance plasmid ($Kan^R$) | This study |
| pCsb2 Y14A | Csb2/I-G with alanine mutation at Y14 amino acid, encoding N-terminal Strep-II tagged protein expressed using p1R plasmid. $Kan^R$ | This study |
| pCsb2 E24A | Csb2/I-G with alanine mutation at E24 amino acid, encoding N-terminal Strep-II tagged protein expressed using p1R plasmid. $Kan^R$ | This study |
| pCsb2 R31A | Csb2/I-G with alanine mutation at R31 amino acid, encoding N-terminal Strep-II tagged protein expressed using p1R plasmid. $Kan^R$ | This study |
| pCsb2 E65A | Csb2/I-G with alanine mutation at E65 amino acid, encoding N-terminal Strep-II tagged protein expressed using p1R plasmid. $Kan^R$ | This study |
| pCsb2 Y90A | Csb2/I-G with alanine mutation at Y90 amino acid, encoding N-terminal Strep-II tagged protein expressed using p1R plasmid. $Kan^R$ | This study |
| pCsb2 Y236A | Csb2/I-G with alanine mutation at Y236 amino acid, encoding N-terminal Strep-II tagged protein expressed using p1R plasmid. $Kan^R$ | This study |
| pCsb2 E410A | Csb2/I-G with alanine mutation at E410 amino acid, encoding N-terminal Strep-II tagged protein expressed using p1R plasmid. $Kan^R$ | This study |
| pCsb2 H520A | Csb2/I-G with alanine mutation at H520 amino acid, encoding N-terminal Strep-II tagged protein expressed using p1R plasmid. $Kan^R$ | This study |
| pCsb2 D530A | Csb2/I-G with alanine mutation at D530 amino acid, encoding N-terminal Strep-II tagged protein expressed using p1R plasmid. $Kan^R$ | This study |
| pCsb2 ΔC | Cas5-like domain of Csb2/I-G without the C-terminal regions (from 251 – 545 amino acids), encoding N-terminal Strep-II tagged protein expressed using p1R plasmid. $Kan^R$ | This study |

| | | |
|---|---|---|
| pCsb2 ΔN | Cas6-like domain of Csb2/I-G without the N-terminal regions (from 1– 250 amino acids), encoding N-terminal Strep-II tagged protein expressed using p1R plasmid. $Kan^R$ | This study |
| pCsb1/I-G | *csb1* gene from *B. animalis* encoding N-terminal Strep tagged protein inserted in p13S-R plasmid. Spectinomycin resistance plasmid ($Spc^R$) | This study |
| pCsb1 E78A | Csb1/I-G with alanine mutation at E78 amino acid, encoding N-terminal Strep-II tagged protein expressed using p13S-R plasmid. $Spc^R$ | This study |
| pCsb1 H122A | Csb1/I-G with alanine mutation at H122 amino acid, encoding N-terminal Strep-II tagged protein expressed using p13S-R plasmid. $Spc^R$ | This study |
| pCsb1 R123A | Csb1/I-G with alanine mutation at R123 amino acid, encoding N-terminal Strep-II tagged protein expressed using p13S-R plasmid. $Spc^R$ | This study |
| pCsb1 R179A | Csb1/I-G with alanine mutation at R179 amino acid, encoding N-terminal Strep-II tagged protein expressed using p13S-R plasmid. $Spc^R$ | This study |
| pCsb1 R320A | Csb1/I-G with alanine mutation at R320 amino acid, encoding N-terminal Strep-II tagged protein expressed using p13S-R plasmid. $Spc^R$ | This study |
| pCsb3/I-G | *csb3* gene from *B. animalis* encoding N-terminal 6xHis tagged protein inserted in pQE2 plasmid. Ampilcilin resistance plasmid ($Amp^R$) | This study |
| pCascade/I-G | Cascade operon (Csb2-Csb1-Csb3) encoding N-terminal 6xHis tagged protein inserted in pQE2 plasmid. $Amp^R$ | This study |
| pC1H_Cascade | Cascade operon (Csb2-Csb1-Csb3) with mutation at Csb1 H122 amino acid, encoding N-terminal 6xHis tagged protein inserted in pQE2 plasmid. $Amp^R$ | This study |
| pC1R_Cascade | Cascade operon (Csb2-Csb1-Csb3) with mutation at Csb1 R123 amino acid, encoding N-terminal 6xHis tagged protein inserted in pQE2 plasmid. $Amp^R$ | This study |
| pC2H_Cascade | Cascade operon (Csb2-Csb1-Csb3) with mutation at Csb2 H520 amino acid, encoding N-terminal 6xHis tagged protein inserted in pQE2 plasmid. $Amp^R$ | This study |
| pC1H_C2H Cascade | Cascade operon (Csb2-Csb1-Csb3) with mutation at Csb1 H122 and Csb2 H520 amino acids, encoding N-terminal 6xHis tagged protein inserted in pQE2 plasmid. $Amp^R$ | This study |
| pC1R_C2H Cascade | Cascade operon (Csb2-Csb1-Csb3) with mutation at Csb1 R123 and Csb2 H520 amino acids, encoding N-terminal 6xHis tagged protein inserted in pQE2 plasmid. $Amp^R$ | This study |
| pCR_Array/I-G | I-G CRISPR array harbouring 7 repeat units interspersed by 6 identical spacer units from *B. animalis*, inserted in 13S-R plasmid, $Spc^R$ | This study |
| pCas3/I-G | *cas3* gene from *B. animalis* encoding N-terminal Strep tagged protein inserted in p1R plasmid. Kanamycin resistance plasmid ($Kan^R$) | This study |
| pT/I-G | Target DNA sequence with TTT PAM inserted in 13S-R plasmid. $Spc^R$ | This study |
